# Acute in vivo proximity labeling for membrane targeted proteomics in neuronal circuits

**DOI:** 10.64898/2025.12.04.691871

**Authors:** Maribel Anguiano, Run Zhang, Melanie Robles, Kaden P. Adams, Michelle R. Salemi, Brett S. Phinney, Christopher S. Leung, Ethan M. Fenton, Kuldeep Giri, Elinor Lewis, Sophia Lin, Jennifer L. Whistler, Alex S. Nord, Christina K. Kim

## Abstract

Despite the growing use of proximity labeling tools, there are limited techniques available to identify subcellularly localized proteins within targeted neuronal circuits on acute timescales in vivo. We engineered membrane targeted versions of the proximity labeling enzyme TurboID and validated their use for proteomic discovery in the mouse brain. We optimized an in vivo protocol to identify proteins in cell bodies and long range neuronal projections, and proteins differentially detected during acute drug labeling windows.

**SUMMARY:** A major goal within molecular systems neuroscience is to bridge the study of neuronal circuit function with changes in protein expression and localization in awake behaving animals. However, there are limited tools for capturing changes in subcellularly-defined proteomes within neuronal circuits during activity-gated timescales in vivo. Here, we engineered targeted versions of the proximity labeling enzyme TurboID, to tag proteins at the neuronal membrane during a user-delivered biotin injection. We optimized a labeling strategy that enables a one-to-two-hour labeling window and tagged proteins in medial prefrontal cortex (mPFC) cell bodies and corresponding axons in a downstream projection. We performed proteomics to identify proteins enriched in mPFC cell bodies and terminals, and upregulated in mPFC cell bodies following an acute cocaine injection. These advancements enable the detection of proteins at the subcellular level within short labeling windows, allowing identification of stimulus-specific proteomes in behaving mice.

## INTRODUCTION

Membrane proteins play an essential role in neuronal signaling and plasticity, contributing significantly to the regulation of neuronal communication within functional neuronal circuits. Identifying cell-type specific proteins at and around the membrane in the living brain, and at specific timepoints, may be crucial for deeper insights into cellular processes and mechanisms underlying neuronal circuit function. While protein affinity purification techniques^1^, such as affinity pulldown and co-immunoprecipitation, remain a gold standard for identifying protein-protein interactions (PPIs) and protein complexes of specific targets^2,3^, these approaches can be challenging to implement broadly in the living brain; for example, to identify all proteins present at a given subcellular neighborhood of cell-types in a brain region.

Instead, researchers have taken advantage of recent advancements in proximity labeling (PL) techniques to tag and enrich proteins using genetically-encodable and targetable enzymes. Biotinylation-based proximity labeling primarily relies on two categories of engineered enzymes called biotin ligases and peroxidases^4^. The engineered peroxidases, such as APEX^5^, APEX2^6^, APEX3^7^, and horseradish peroxidase (HRP)^8,9^, can catalyze the oxidation of biotin-phenols in the presence of hydrogen peroxide (H_2_O_2_), producing short-lived biotin-phenoxyl radicals^10,11^. These highly reactive radicals are capable of biotinylating electron-rich amino acids within proximal proteins^12^. But despite its rapid temporal resolution, the application of peroxidase-based biotinylation in vivo is limited by the potential cellular toxicity introduced by H_2_O_2_, requiring the actual labeling of proteins in an ex vivo preparation^13-15^ (**Table 1**). APEX2 has nonetheless been employed to identity cell-type specific proteomes in both the nucleus and at the membrane of mouse neurons of the striatum^13^; and in cell bodies and axon terminals of dopamine neurons^14^.

**Table 1.** Labeling features of proximity labeling tools used for proteomics in mouse brain tissue.

| Enzyme | Required substrate | In vivo compatibility | Subcellular targeting | Reported labeling window | References |
| --- | --- | --- | --- | --- | --- |
| APEX2 | Biotin phenol + H<sub>2</sub>O<sub>2</sub> | Toxic, labeling performed ex vivo | Cytosol, nucleus, membrane | 1 hour biotin phenol, 1 min H<sub>2</sub>O<sub>2</sub> | Dumrongprechachan et al., 2021<sup>13</sup> |
|  |  |  | Membrane axon terminals | 1 hour biotin phenol, 3 min H<sub>2</sub>O<sub>2</sub> | Hobson et al., 2022<sup>14</sup> |
| HRP | Biotin phenol + H<sub>2</sub>O<sub>2</sub> | Toxic, labeling performed ex vivo | Soma, synapse | 1 hour biotin phenol, 3 min H<sub>2</sub>O<sub>2</sub> | Shuster et al., 2023<sup>15</sup> |
| iBioID | Biotin | Non-toxic, labeling performed in vivo | Soma, Synapse | 7 days of i.p. biotin injections | Uezu et al, 2016<sup>17</sup> |
| TurboID | Biotin | Non-toxic, labeling performed in vivo | Cytosol | 2 weeks of biotin in water | Rayaprolu et al., 2022<sup>24</sup> |
| Split-TurboID | Biotin | Non-toxic, labeling performed in vivo | Astrocyte-neuron junctions | 7 days of s.c. biotin injections | Takano et al., 2020<sup>21</sup> |

In contrast, the engineered biotin ligases BioID^16,17^ and TurboID^18^ are PL tools that catalyze the formation of reactive biotin-5’-AMP using ATP and biotin, without the need of H_2_O_2_. This reactive biotin-5’-AMP molecule diffuses to label lysine residues within proximal proteins^19^, enabling proteomic mapping with spatial precision in living cells. Both BioID and TurboID have been used in vivo in mice to identify proteins present at synapses^17,20-22^ or throughout the cytosol of different neuronal cell-types^23,24^. For example, the first transgenic TurboID mouse strain demonstrated the identification of 2,143 proteins present in neuronal cell-types across the brain^24^. However, neither variant has been adapted thus far to identify intracellular membrane proteins in vivo. Furthermore, prior studies also required multiple days of biotin induction via repeated intraperitoneal injections^17,20,22,25^, due to the lower efficiency of BioID (**Table 1**). This multi-day labeling requirement precludes the ability to capture acute changes in membrane-associated proteins in response to specific events in vivo. While TurboID is reported to be more catalytically active compared to BioID, producing more of the reactive biotin-5’-AMP substrate per unit of time, most studies in vivo have still relied on 5-7 days of biotin injections for labeling^23,24^.

Thus, to enable the investigation of dynamic membrane proteomic changes that may occur in response to acute stimuli, we characterized a new membrane-targeted TurboID, Nrxn-TurboID, that can be used in vivo and demonstrated its ability to robustly biotinylate proteins after just a single session of biotin administration. We applied Nrxn-TurboID to identify proteins present in mouse medial prefrontal cortex (mPFC) cell bodies, and in two of its downstream axonal projections to the nucleus accumbens (NAc) and the periaqueductal grey (PAG)^26-28^. We also demonstrated that virally transduced Nrxn-TurboID can be used to identify acute molecular changes occurring within mPFC cell bodies during a 1-hour window following a cocaine injection. Our study provides a robust platform to perform membrane-targeted proteomics across different subcellular locations in vivo, requiring only a single labeling session within individual mice to collect comprehensive proteomic maps of neuronal cell bodies and terminals.

## RESULTS

### Design and engineering of potential membrane-TurboID variants

We generated various membrane-TurboID candidates by tethering the enzyme to different membrane-targeting motifs, and then compared them to a cytosolic TurboID (**Fig. 1A**). We chose several different engineered membrane-scaffolding motifs that have been previously demonstrated in the literature to target multi-component heterologous protein fusions to neuronal membranes in culture and/or in vivo in the mouse: a truncated Neurexin-3β-derived transmembrane protein^29^ (abbreviated here as Nrxn), which is a minimal membrane-trafficking variant of the endogenously occurring pre-synaptic protein, Neurexin; Nrxn fused to the CIBN^30^ protein (a truncated version of the natural CIB1 cryptochrome-interacting basic-helix-loop-helix 1 protein), which is an unrelated synthetic light-modulated protein that has nevertheless been shown to enhance the membrane trafficking of Nrxn-fused constructs in neurons^29,31^; an inert, truncated version of the human T cell protein CD4^32^, which has been used to target synthetic proteins to the membrane; and the more canonically described CAAX prenylation motifs^33^ (C=cysteine, A=aliphatic amino acid, X=terminal amino acid), which trigger a series of posttranslational modifications to induce translocation to the membrane. Note that these targeting motifs are intended to localize TurboID to the general vicinity of the intracellular membrane, rather than to serve as specific “bait” or defined “protein interaction partner” for downstream proteomics.

**Figure 1.**
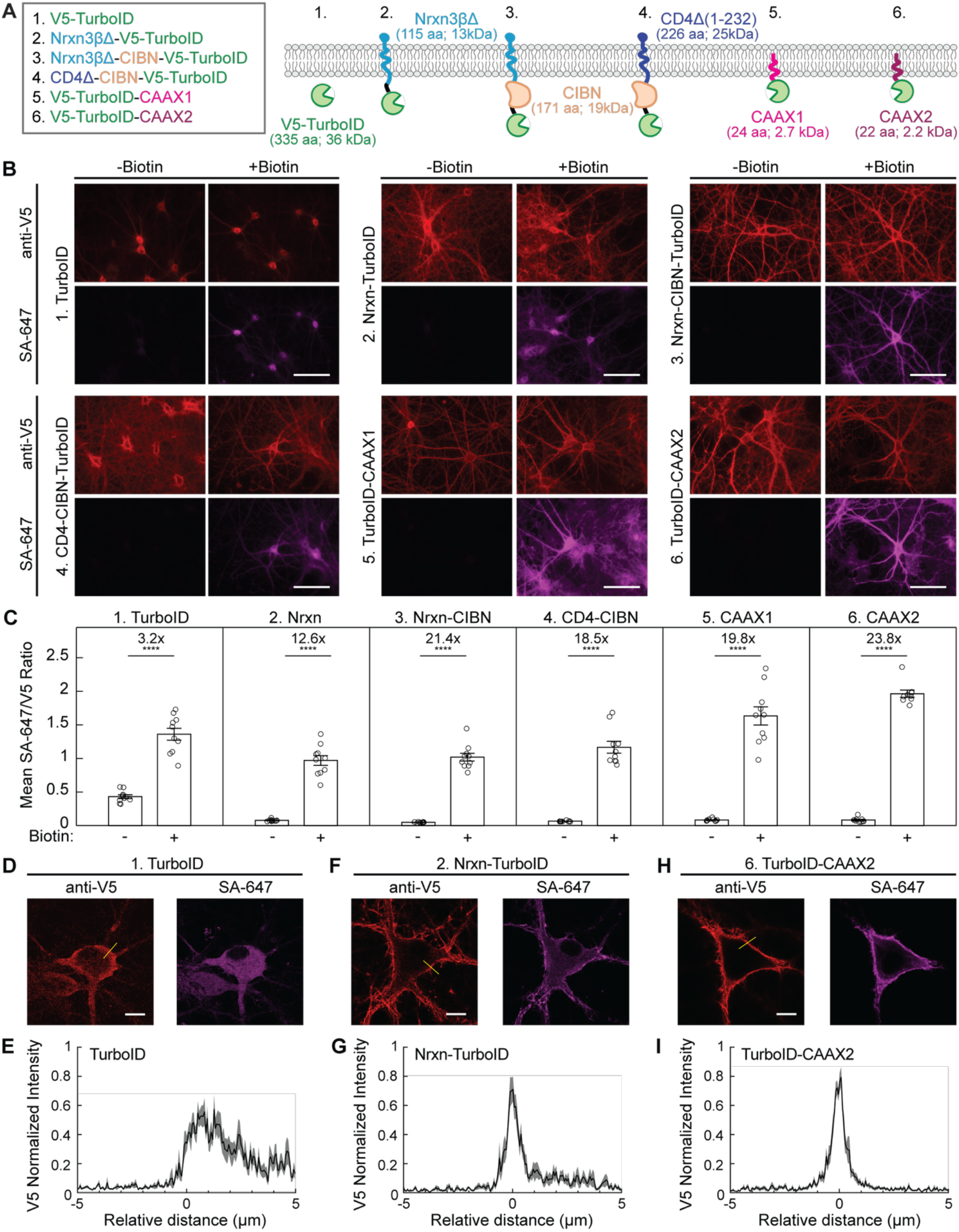
Design and validation of different membrane-TurboID variants. **(A)** Schematic depicting two general strategies for targeting TurboID to the neuronal membrane. Middle: the N terminus of TurboID was fused to the C terminus of a truncated Neurexin3β (Nrxn), Nrxn-CIBN, or CD4-CIBN. Right: the C terminus of TurboID was modified with two different CAAX signaling motifs. **(B)** Example 40x images of anti-V5 and SA-647 staining of cultured neurons expressing 1 of 6 TurboID variants. Neurons were treated with either vehicle or 50 µM biotin for 10 minutes, then fixed and stained for TurboID localization and biotinylation. Scale bar, 100 µm. **(C)** Quantification of the mean SA-647 divided by anti-V5 cell fluorescence across the entire FOV of data from panel B (*n* = 9-10 FOV per condition; 2-tailed Mann-Whitney U test, ****p=0.0001). **(D)** Example 100x images of anti-V5 and SA-647 staining of cultured neurons expressing cytosolic TurboID and treated with biotin as in panel B. The yellow line indicates pixels used to quantify TurboID membrane localization for the summary graph in panel E. Scale bar, 10 µm. **(E)** Quantification of the mean anti-V5 membrane localization of TurboID across neurons (*n* = 10 neurons, data plotted as mean +/-standard error of the mean (SEM)). **(F-I)** Same as panels (D-E), except for cultured neurons infected with Nrxn-TurboID (**F-G**) and TurboID-CAAX2 (**H-I**). See also **Figure S1**.

**Figure 1A** shows schematics of all constructs we generated, using different combination of the above-described membrane-targeting motifs: Nrxn-TurboID, Nrxn-CIBN-TurboID, CD4-CIBN-TurboID, TurboID CAAX1 (from the K-Ras GTPase), and TurboID-CAAX2 (from the H-Ras GTPase). A V5 epitope tag was included at the N-terminus of TurboID in all constructs for immunohistochemistry of TurboID targeting.

To evaluate the extent of biotin-dependent protein labeling of these variants in cultured neurons, we performed fluorescence imaging and quantification across multiple fields of view (FOVs) in membrane-TurboID expressing cells treated with or without biotin (+/-biotin) and stained for TurboID expression (anti-V5) and biotinylated proteins using fluorescently tagged Streptavidin (SA-647). Fluorescence imaging confirmed strong biotinylation signal upon treatment for all tested membrane-TurboID variants (**Fig. 1B,C**). All variants also showed strong expression and biotinylation in the neuronal processes, compared to the cytosolic TurboID, indicating an increased membrane localization of these new variants.

We proceeded with two different variants for in vivo testing: Nrxn-TurboID and TurboID-CAAX2. High-magnification imaging and line-scanning analysis showed clear membrane localization of these variants in comparison to cytosolic TurboID (**Fig. 1D-I**). While Nrxn-CIBN-TurboID and CD4-CIBN-TurboID also showed robust membrane targeting (**Fig. S1A-D**), we chose Nrxn-TurboID because it was smaller in size—potentially resulting in less steric hindrance for labeling proteins nearby the membrane. We chose the TurboID-CAAX2 variant because we observed that TurboID-CAAX1 resulted in biotinylated protein signal throughout the entire neuron, not only at the membrane (**Fig. S1E-F**). For subsequent in vivo testing experiments, we refer to the CAAX2 variant as TurboID-CAAX.

### Rapid proximity labeling in vivo using membrane-targeted TurboID variants

To test our two membrane-TurboID variants in vivo, we injected either AAV2/1-Synapsin-Nrxn-TurboID or AAV2/1-Synapsin-TurboID-CAAX in the mPFC of mice (**Fig. 2A**; see **Fig. S2** for representative viral spread and targeting). The AAV2/1 serotype and synapsin promoter enables infection of broad neuronal cell-types across multiple brain regions. Given that prior in vivo proximity labeling studies using BioID or TurboID have been performed over multiple days of biotin labeling **(Table 1)**, we asked whether our TurboID variants and AAV approach could be used for acute proximity labeling in vivo over the span of hours, instead of days. For biotin labeling, we gave mice two intraperitoneal (IP) injections of biotin, 1 hour apart, and then sacrificed and perfused them with ice-cold phosphate buffered saline (PBS) followed by 4% paraformaldehyde (PFA) 1 hour later. TurboID expression and the extent of biotinylation were visualized by staining with anti-V5 primary antibody and SA-647, respectively. For both Nrxn-TurboID (**Fig. 2B-C**) and TurboID-CAAX (**Fig. 2D-E**), we found that a two-hour biotin labeling window was sufficient to see a significant level of protein labeling in the mPFC cell bodies in biotin treated conditions compared to negative control conditions.

**Figure 2.**
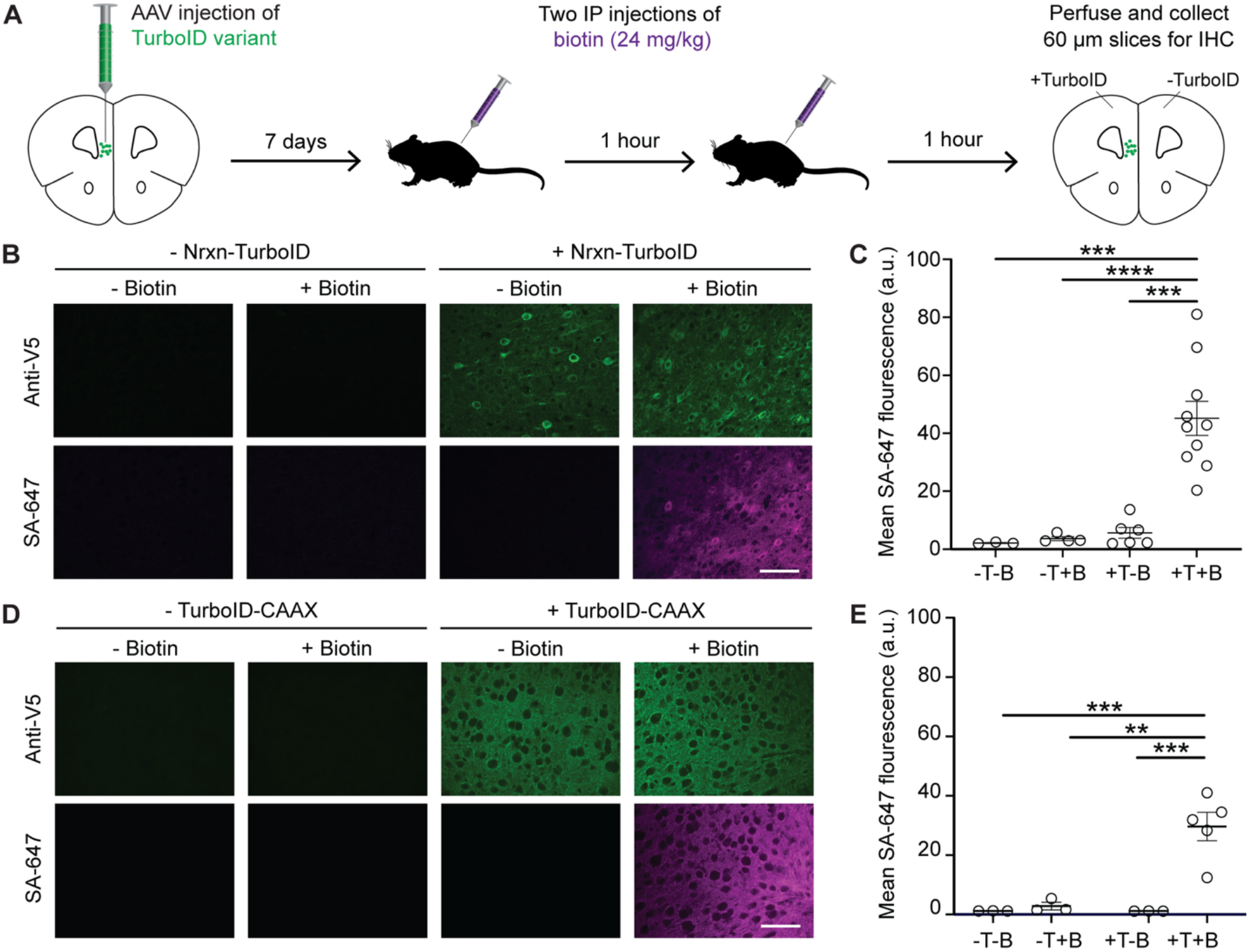
Rapid in vivo cell body labeling using different TurboID variants. **(A)** Schematic depicting experimental protocol. Mice were injected with AAV2/1-Syn-Nrxn-TurboID or AAV2/1-Syn-TurboID-CAAX in the mPFC, then one week later they were injected with two IP biotin (24 mg/kg) or vehicle injections, one hour apart. The brains were then collected for immunohistochemistry (IHC). **(B)** 40x images of anti-V5 and SA-647 staining of mPFC neurons from mice expressing Nrxn-TurboID and treated with biotin as in panel A or treated with vehicle. Scale bar, 75 µm. **(C)** Quantification of mean SA-647 fluorescence from experiment in panel B (*n* = 3-10 FOVs per condition; 2-way ANOVA with significant interaction F_(1,19)_=9.994, p=0.0051; followed by Tukey’s multiple comparison test ***p<0.001, ****p<0.0001). Scale bar, 75 µm. **(D)** Same as panel B, except for mice expressing TurboID-CAAX in mPFC. **(E)** Quantification of mean SA-647 fluorescence from experiment in panel D (*n* = 3-5 FOVs per condition; 2-way ANOVA with significant interaction, F_(1,10)_=12.88, p=0.0049; followed by Tukey’s multiple comparison test **p<0.01, ***p<0.001). See also **Figure S2**.

### Proteomics of membrane-associated proteins in mPFC neurons in vivo using Nrxn-TurboID

Next, we validated the neuronal specificity of our TurboID variants at the level of the proteome, starting with the Nrxn-TurboID construct. We followed the same surgical and labeling timeline as in Figure 2, but then microdissected the mPFC following an ice-cold PBS-only perfusion (**Fig. 3A**). After microdissection, we homogenized and lysed the tissue and then extracted the labeled proteins using streptavidin magnetic bead enrichment (**Fig. 3B**). We confirmed the enrichment of biotinylated proteins by Western blot, finding stronger SA-HRP labeling across multiple protein sizes in samples from mice expressing Nrxn-TurboID and injected with biotin, compared to samples from mice expressing Nrxn-TurboID but injected with vehicle instead of biotin (**Fig. 3C, top, Fig. S3A-C**). TurboID itself (anti-V5) was also enriched in samples from biotin-treated compared to untreated mice, since the enzyme undergoes auto-labeling in the presence of biotin (**Fig. 3C, bottom**). After quality control of the samples, we performed on-bead digestion of proteins and LC-MS/MS analysis using data-independent acquisition (DIA) proteomics (**Fig. S4A-D**). As expected, the Nrxn-TurboID construct captured a substantial number of proteins in +biotin samples compared to negative control -biotin samples, validating that our protocol is properly isolating biotinylated proteins above endogenous background levels. We filtered an initial list of mPFC candidates using the following criteria: they must be detected with 2 or more unique peptides and have a +/-Biotin fold change ≥ 1.5 (L2FC ≥ 0.585) with a Q-value < 0.05 (**Fig. 3D**). We also removed known universal human and mouse contaminants detected in mass spectrometry data (**Table S4**). This resulted in 1,881 proteins detected in the Nrxn-TurboID dataset (**Fig. 3E**). The min-max normalized protein expression across biological replicates is shown for the top 20 candidates with the largest significant +/-Biotin fold change (**Fig. 3F**).

**Figure 3.**
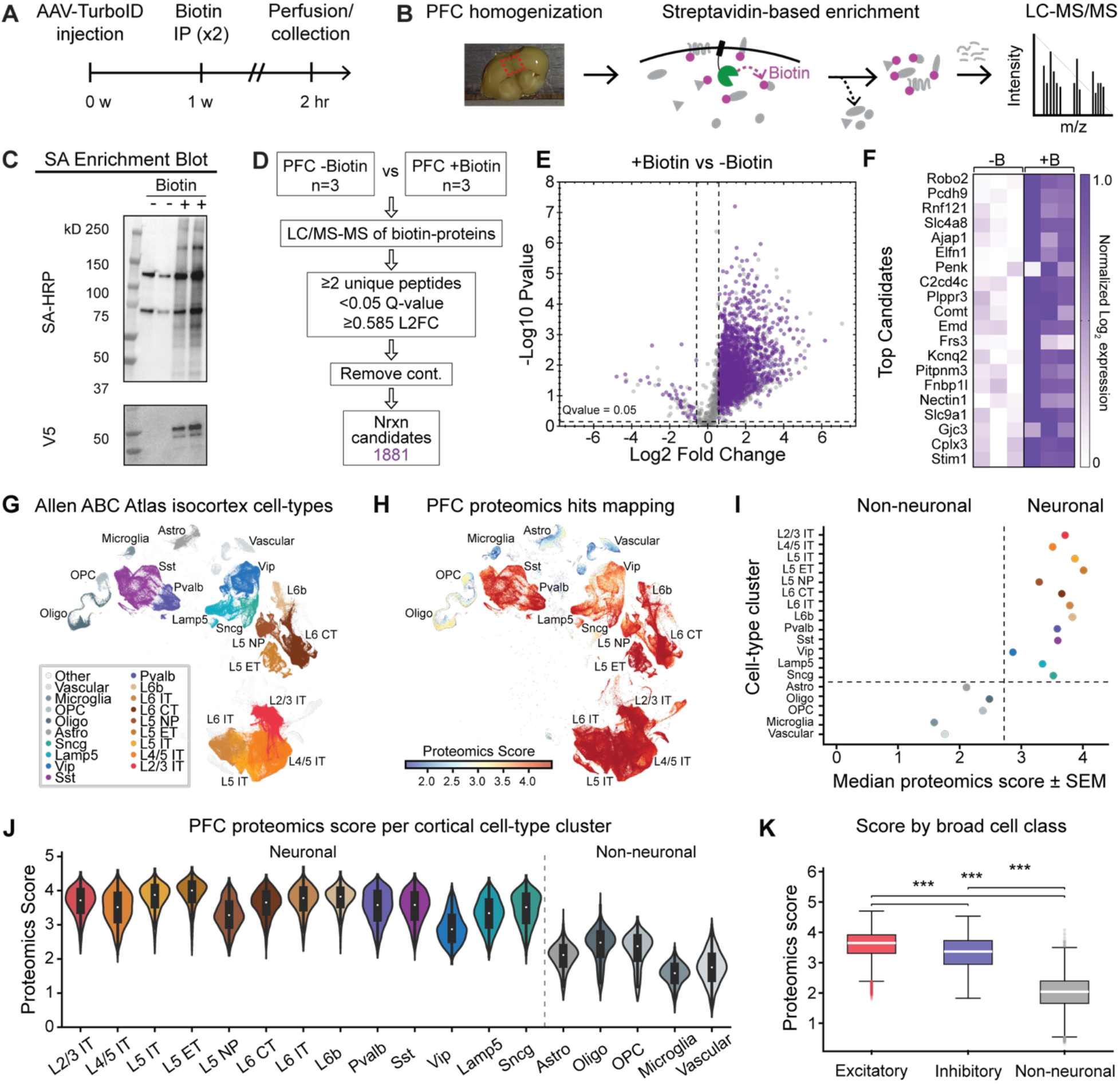
In vivo protein labeling and proteomics of mPFC neurons using Nrxn-TurboID. **(A-B)** Schematics for TurboID experiments. AAV2/1-Syn-Nrxn-TurboID was bilaterally injected into mPFC. 1 week later, mice were given two IP biotin (24 mg/kg) or vehicle injections, one hour apart. Mice were flushed with PBS through transcranial perfusion, and then brains were collected for mPFC microdissection. Protein extraction was performed, followed by streptavidin magnetic bead enrichment. After on-bead digestion, peptides were analyzed using DIA proteomics. **(C)** Western blot showing SA-HRP and anti-V5 staining of biotin-enriched samples. **(D)** Workflow for data collection and filtering steps. We performed three biological replicates per experimental condition. Proteins were filtered and identified as candidates if they were detected with ≥ 2 unique peptides and had a +/-Biotin L2FC ≥ 0.585 (1.5) with a Q-value < 0.05. **(E)** Volcano plot showing results of +/-biotin differential protein expression analysis for Nrxn-TurboID. **(F)** Heatmap of min-max normalized log_2_ expression data for the top 20 candidates with the largest +/-Biotin fold change in the Nrxn-TurboID mPFC dataset. **(G)** UMAP showing cell type clusters from the Allen ABC Atlas from the mouse isocortex. **(H)** Each cell in the UMAP pseudocolored based on a “Proteomics Score’ calculated from the list of mPFC protein candidates from panel D (see methods). **(I)** Median proteomics scores calculated per cell-type cluster (Non-neuronal and Neuronal). **(J)** Violin plots showing distribution of Proteomics Scores across neuronal and non-neuronal clusters. (**K**) Distribution of proteomics scores pooled across broad excitatory neuronal cell-types, inhibitory neuronal cell-types, and non-neuronal cell-types. Mann-Whitney U-test with Bonferroni correction ***adj. p<0.001. Data plotted as median +/-SEM. See also **Figures S3 and S4**.

Given the immense diversity of cell-types and tissue in the mouse brain, we first asked whether our enrichment protocol of biotinylated proteins is properly resulting in the detection of proteins present in neurons vs non-neuronal cells. We took our list of mPFC protein candidates and compared them to the Allen ABC Atlas of mouse single-cell RNA sequencing data from the isocortex (**Fig. 3G**). We created a proteomics score for each cell in the ABC atlas, comparing the similarity of the detected RNA transcripts in that cell with the bulk proteomics data from our Nrxn-TurboID mPFC dataset (**Fig. 3H**). We calculated the median proteomics score within each cell-type cluster, and found that neuronal cell-types had a greater mapped proteomics score compared to non-neuronal cell-types (**Fig. 3I,J**). Both the broad classes of excitatory and inhibitory neurons had a greater mapped proteomics score compared to that of non-neuronal cell-types (**Fig. 3K**).

Next, we aimed to determine the subcellular specificity of the mPFC Nrxn-TurboID proteome. Ideally, in proximity labeling experiments, we would compare our list of candidate proteins to a known “true positive list” and “false positive list” of proteins that should be either represented or excluded from our proteome. This allows us to determine a L2FC cut-off that results in a false discovery rate (FDR) of <0.10, and also determine the sensitivity and specificity of our proteome. However, unlike proximity labeling proteomic datasets collected in uniform cultured cell-types (HEK293T cells^18^), there are limited databases for obtaining a “ground truth” list of true positive membrane proteins that are present in a given sub-type of neuron. Thus as our true positive list, we used a dataset of 57 identified somatic neuronal membrane proteins, detected in a study performing proximity labeling in mouse striatal neurons using a membrane-targeted APEX2 (LCK-APEX2^13^). As our false positive list, we curated a list of 62 nuclear proteins starting from the same study (H2B-APEX2). 45 of the “true positive” proteins and 39 of the “false positive” proteins were detected in our list of Nrxn-TurboID candidates (**Fig. 4A**). Receiver operating characteristic (ROC) curve analysis showed that our full candidate list had an area under the curve (AUC) of 0.728 (**Fig. 4B**). To achieve an FDR<0.1, a L2FC cutoff of 2.259 was defined (**Fig. 4C,D**). Using this new threshold, a refined list of 280 mPFC membrane candidates was determined (**Fig. 4E**). A gene ontology (GO) enrichment analysis using ShinyGO^34^ identified the top 10 associated cellular components for the 280 candidates, all showing membrane specificity, including “cell junction”, “synapse”, “cell projection”, “neuron projection” (**Fig. 4F**).

**Figure 4.**
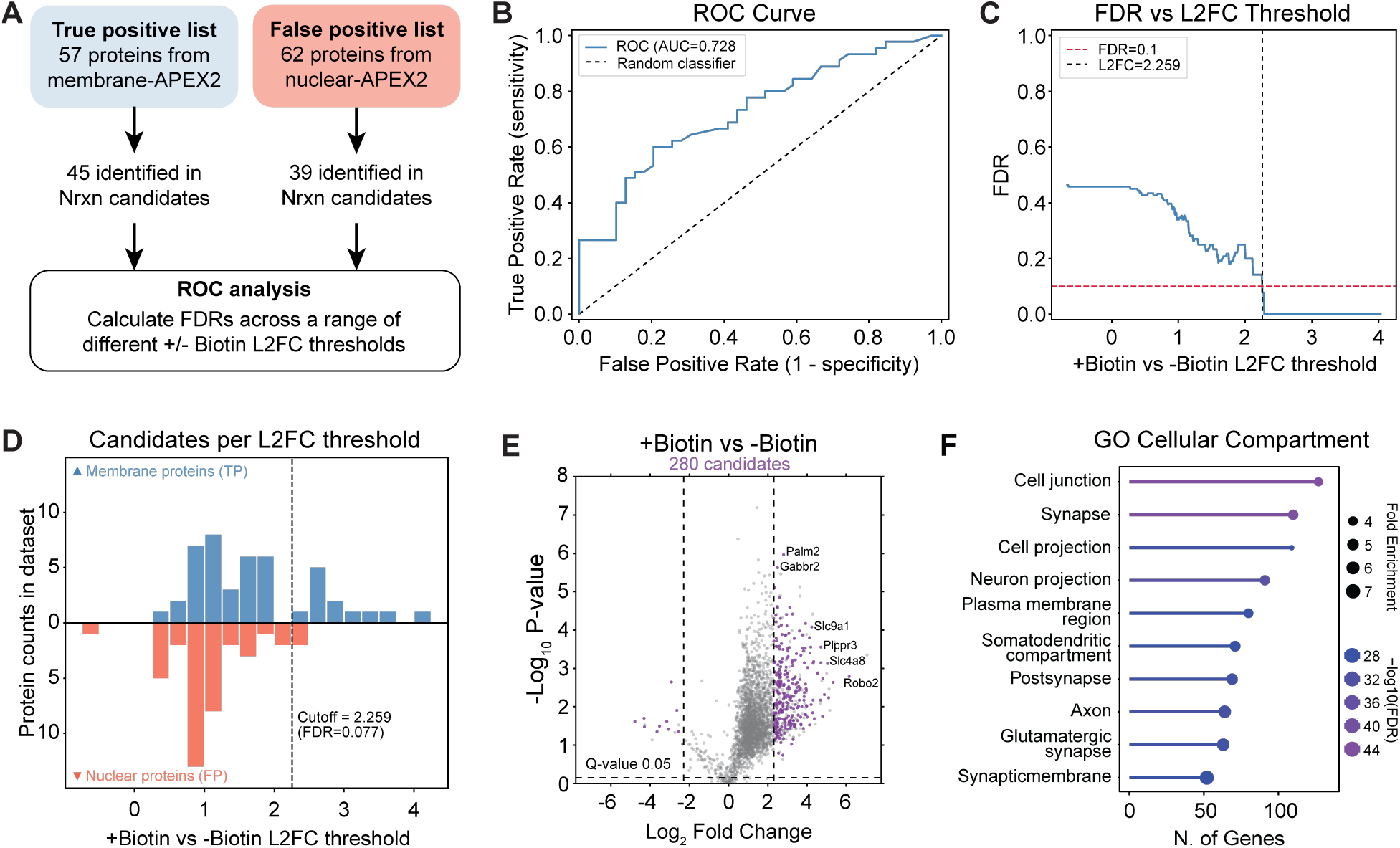
ROC analysis of mPFC Nrxn-TurboID candidates. **(A)** Schematic depicting strategy for ROC analysis. A curated TP list of neuronal membrane proteins was built from a published LCK-APEX2 mouse dataset^13^, and a curated FP list of nuclear membrane proteins was taken from a published H2B-APEX2 mouse dataset^13^ and UniProtKB searches. **(B)** ROC curve showing TP rate and FP rate determined using different L2FC thresholds. **(C)** Plot of the FDR versus +/-Biotin L2FC threshold, with the dashed vertical line indicating the L2FC at which the FDR<0.1. **(D)** Histogram showing the number of TP (blue) and FP (red) proteins present at different +/-Biotin L2FC thresholds. **(E)** Volcano plot with a more stringent L2FC threshold resulting in an FDR<0.1. **(F)** Top 10 enriched GO Cellular Compartments determined from the 280 filtered candidates in panel (E). See also **Table S5**.

### Proteomics of membrane-associated proteins in mPFC neurons in vivo using TurboID-CAAX

We next performed another enrichment experiment using TurboID-CAAX expressed in the mPFC. Tissue and proteins were processed and enriched exactly as described in **Fig. 3**. As with the Nrxn-TurboID construct, we confirmed enrichment of biotinylated proteins by Western blot (**Fig. S3D-F**). However, overall, the TurboID-CAAX dataset detected fewer peptides than the Nrxn-TurboID dataset (5,488 unique peptides versus 18,455 unique peptides; **Fig. S4E-H**). We performed the same initial filtering steps to identify 508 potential candidates (**Fig. S5A-D**). Here, a smaller subset of 22 true positive and 11 false positive proteins were identified in the CAAX candidates, and ROC curve analysis found an AUC of 0.864 and a L2FC cutoff of 1.155 to achieve an FDR<0.1 (**Fig. S5E-H**). GO cellular component analysis of the remaining 387 filtered candidates also showed membrane specificity (**Fig. S5I**).

We compared the full 1,881 Nrxn candidate list with the full 508 CAAX candidate list, and found 423 proteins shared between the lists, representing 22% of all Nrxn candidates and 83% of all CAAX candidates (**Fig. S6A**). We then investigated the “exclusive” candidates that were found only in either the Nrxn-TurboID or TurboID-CAAX datasets, and asked why they were not present in both datasets. We searched for the 1,458 “Nrxn-only” candidates within the unfiltered list of identified CAAX proteins, to determine why they were excluded as CAAX candidates. We found that 61% of “Nrxn-only” candidates were not detected at the peptide level in the TurboID-CAAX LC-MS/MS runs (**Fig. S6B top**, grey); 27% were detected in the CAAX runs but did not have a significant Q-value comparing +biotin vs -biotin conditions (**Fig. S6B**, yellow; Q-value ≥ 0.05); 2% were significantly downregulated in the +biotin vs -biotin conditions in the CAAX dataset (**Fig. S6B**, orange); and <1% were upregulated in the +biotin vs -biotin CAAX dataset but fell below our defined threshold of a 1.5-fold enrichment (**Fig. S6B**, purple). Finally, 10% of the Nrxn-only candidates reached all other filtering criteria in the CAAX runs but were detected with < 2 peptides (**Fig. S6B**, blue). Similarly, we searched for the 85 “CAAX-only” candidates in the total unfiltered Nrxn-TurboID dataset, and found that 52% of the CAAX-only candidates were not detected in the Nrxn dataset; 7% were downregulated in +biotin vs -biotin conditions; 25% were enriched in +biotin vs -biotin conditions but did not meet the 1.5-fold change criteria; and 16% met all other inclusion criteria, but were detected with < 2 peptides (**Fig. S6C**). Using our true positive and false positive lists, we then calculated the estimated specificity and sensitivity for the Nrxn-TurboID and TurboID-CAAX datasets dependent on proteome size (using decreasing L2FC cutoffs; **Fig. S6D,E**). Based on this comparison, the Nrxn proteome maintains a specificity >90% up to a proteome size of 679 candidates, while the CAAX proteome maintains specificity >90% up to 451 candidates. At these proteome sizes, the Nrxn proteome has a sensitivity of 39%, while CAAX proteome has a sensitivity of 35%. These data suggest that both variants detect many of the same membrane associated proteins, but that the Nrxn-TurboID has greater sensitivity and more proteins detected while maintaining an appropriate tolerance level of specificity. Thus, we decided to further characterize the Nrxn-TurboID construct to ask how it performs in contexts with sparse protein labeling (e.g. in axon terminals) or in contexts detecting protein localization changes (e.g. in response to drug).

### Subcellularly targeted proteomics at long range mPFC axonal terminals in subcortical regions

We asked whether Nrxn-TurboID enables protein enrichment at axonal terminals of long-range mPFC projections to distal subcortical regions. We chose to enrich axonal mPFC proteins projecting to the medial shell nucleus accumbens (NAc), which is a major cortical projection pathway that is known to regulate reward-seeking behavior^35^. Following the same experiment in Figure 2, mice injected with AAV-Syn-Nrxn-TurboID in mPFC were injected with biotin or saline over 2 hours and then sacrificed and perfused with ice-cold PBS and PFA (**Fig. 5A**). Coronal sections from the NAc were stained and imaged for anti-V5 TurboID expression and SA-647 biotinylation in mPFC axons (**Fig. 5B-C**). Even in axon terminals, we were able to detect elevated SA-647 labeling in biotin-treated mice expressing Nrxn-TurboID. In contrast, mice injected with TurboID-CAAX in mPFC did not exhibit a robust +/-biotin fold-change in SA-647 labeling in mPFC axon terminals in medial shell NAc (**Fig. S7**). This could be because the Nrxn fusion protein, even though a truncated form, promotes better trafficking to pre-synaptic axon terminals compared to the CAAX; or because of the overall lower sensitivity of the TurboID-CAAX compared to Nrxn-TurboID.

**Figure 5.**
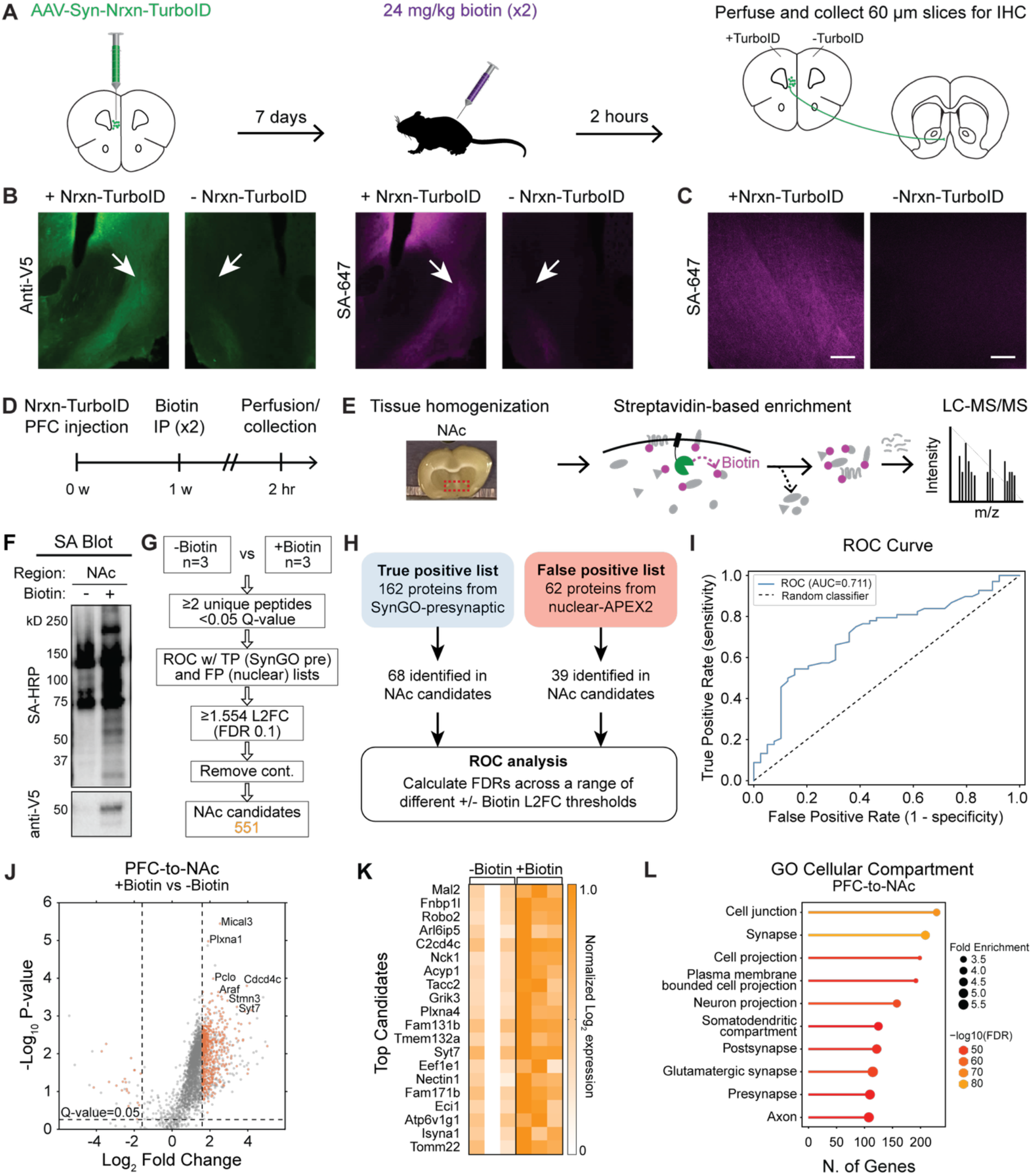
In vivo protein labeling and proteomics of mPFC-to-NAc neuron terminals using Nrxn-TurboID. **(A)** Schematic depicting experimental protocol. Mice were injected unilaterally in the mPFC with AAV2/1-Syn-Nrxn-V5-TurboID. After 1 week, they were injected with two IP biotin (24 mg/kg) or vehicle injections, one hour apart. The brains were then collected for IHC. **(B)** 10x images of the injected and uninjected hemispheres from same mouse, showing the NAc with anti-v5 and SA-647 staining. White arrows point to regions microdissected for proteomics. **(C)** Confocal images of SA-647 staining in NAc. Scale bar, 20 µm. **(D-E)** Schematics for TurboID experiments. AAV2/1-Syn-Nrxn-TurboID was bilaterally injected into mPFC. 1 week later, mice were given two IP biotin (24 mg/kg) or saline injections, one hour apart. Mice were flushed with PBS through transcranial perfusion, and then brains were collected for NAc microdissection. Protein extraction was performed, followed by streptavidin magnetic bead enrichment. After on-bead digestion, peptides were analyzed using DIA proteomics. **(F)** Western blot showing SA-HRP and anti-V5 staining of biotin-enriched samples. **(G)** Workflow for data collection and filtering steps. **(H)** Schematic depicting strategy for ROC analysis. TP list was taken from SynGO presynaptic terms, and the same FP list was used as in Figure 4. **(I)** ROC curve showing TP rate and FP rate determined using different L2FC thresholds. **(J)** Volcano plots for comparison of +/-Biotin protein expression for mPFC terminals in NAc, using the L2FC cutoff with FDR=0.1. **(K)** Heatmap of min-max normalized log_2_ expression data for the top 20 candidates with the largest +/-Biotin fold change. **(L)** Top 10 GO enrichment analysis terms for cellular compartment performed using the filtered protein candidates in NAc. See also **Figures S8 and S9**.

In separate animals following the same experiment in Figure 3, we microdissected and processed the NAc for proteomics in mice expressing AAV-Syn-Nrxn-TurboID in mPFC (**Fig. 5D,E**). After visual confirmation of biotinylated proteins with the enrichment blot (**Fig. 5F, Fig. S3A-C**), we analyzed the enriched proteins from the NAc samples using DIA proteomics (**Fig. S8**). As before, proteins were identified as true candidates if they were detected with 2 or more unique peptides and had a +/-Biotin L2FC ≥ 1.5 with a Q-value < 0.05. Additionally, we excluded common contaminant peptides^36^. This criterion resulted in 1,917 mPFC-to-NAc candidates, that were further narrowed down to a pre-synaptic membrane set of 551 proteins following ROC analysis using a SynGO presynaptic protein true positive list and the same APEX2 nuclear false positive list (**Fig. 5G-K**; **Fig. S9**). The top 10 GO enrichment cellular compartments determined from the 551 candidates in the NAc dataset were associated with membrane and synaptic terms as expected (**Fig. 5L**).

We also asked whether we could detect proteins from mPFC terminals in an even more distal subcortical region, the periaqueductal gray (PAG). This pathway represents another major cortical projection that primarily drives avoidance or aversive behavior^37^.

Histological and Western blot data showed clear enrichment of labeled proteins in Nrxn-TurboID+Biotin treated mice versus negative control conditions (**Fig. S10A-F, Fig. S3A-C**), and we detected an overall enrichment of V5-TurboID in TurboID+Biotin versus TurboID-Biotin samples (**Fig. S11**). However, after filtering, no proteins survived adjusted Q-value thresholding (**Fig. S10G**). This could be because there are fewer mPFC-to-PAG terminals compared to mPFC-to-NAc terminals, or due to lower Nrxn-TurboID expression levels in more distal axonal projections during the limited 1 week viral incubation window. As an exploratory dataset, we analyzed potential mPFC-to-PAG candidates with an unadjusted P-value less than 0.05, resulting in 143 proteins (**Fig. S10H-J**). In this small exploratory dataset, the top candidates also exhibited GO cellular components associated with membrane and synaptic components (**Fig. S10K**).

We compared proteins that were exclusively identified in either the filtered PFC-to-NAc (483 proteins) or exploratory mPFC-to-PAG (82 proteins) candidate proteomes (**Fig. S12A**). We compared the exclusive protein hits to a previously published RNA sequencing (RNA-seq) dataset collected from PFC-to-NAc and PFC-to-PAG neurons^28^. We asked whether exclusive NAc or PAG proteins were also differentially enriched at the RNA level in PFC-to-NAc versus PFC-to-PAG neurons. We first determined all genes that were significantly differentially expressed in either direction between PFC-to-NAc and PFC-to-PAG cells in the RNA-seq dataset (DE genes with L2FC>0.05, adj. p-value <0.05; **Table S6**). We calculated the average L2FC NAc/PAG values for the DE genes that were also found in our exclusive NAc or PAG proteomes (**Fig. S12B**-**C**). The L2FC of NAc/PAG DE genes was >0 for NAc exclusive proteins, and <0 for PAG exclusive proteins; meaning that proteins exclusively found in PFC-to-NAc (but not PFC-to-PAG terminals) correspond to genes that were more enriched in PFC-to-NAc versus PFC-to-PAG neurons at the RNA level, and vice versa.

### Detection of drug-induced changes in the mPFC local membrane proteome

We asked whether the rapid labeling of Nrxn-TurboID would allow the detection of proteomic changes after an acute drug exposure in vivo. Prior BioID and TurboID studies have relied on multiple days of repeated biotin dosing, precluding the ability to capture acute stimulus-induced changes in protein levels in neurons. To test the acute labeling ability of our Nrxn-TurboID AAV-based platform, we asked whether we could detect protein level changes induced near the neuronal membrane of mPFC neurons in response to a drug, such as cocaine. Prior studies have confirmed that cocaine exposure drives physiological changes in mPFC neurons^38,39^, and data suggest that numerous proteins are modulated long-term in mPFC neurons in response to repeated exposure to cocaine^40-42^. Thus, we expressed Nrxn-TurboID in mPFC and administered a single injection of biotin and cocaine and waited for only 1 hour (**Fig. 6A**). We collected mPFC tissue from 3 different conditions: cocaine-biotin (C-B), saline+biotin (S+B), and cocaine+biotin (C+B). Western blot and LC-MS/MS analysis confirmed enrichment of biotinylated proteins above endogenous levels in both S+B and C+B groups (**Fig. 6B, Fig. S13A,B**). We performed two parallel comparisons for analysis, first identifying proteins that were significantly enriched in C+B vs S+B conditions (108 proteins), and then proteins that were enriched in C+B vs C-B (587 proteins; **Fig. 6C, Fig. S14**). The first comparison identified proteins upregulated at the membrane in response to cocaine; the second comparison identified TurboID vs endogenously biotinylated proteins. For the cocaine versus saline comparison, we normalized the total protein counts based on the median quantity of peptides that were identified across all runs, to mitigate differences in tissue dissection size or AAV infection across samples (**Fig. S13C,D**). The Cocaine+Biotin versus Cocaine-Biotin dataset was not normalized, as we expect only background biotinylated proteins in the -Biotin condition (**Fig. S15A**). We compared the overlapping hits between the two datasets and identified 106 proteins that were significantly upregulated by drug (**Fig. 6D**). As expected, most of the filtered candidates found in our cocaine datasets overlapped with the filtered candidates found in the longer, 2-hour labeling Nrxn-TurboID dataset in Figure 3 (**Fig. S15B-D**).

**Figure 6.**
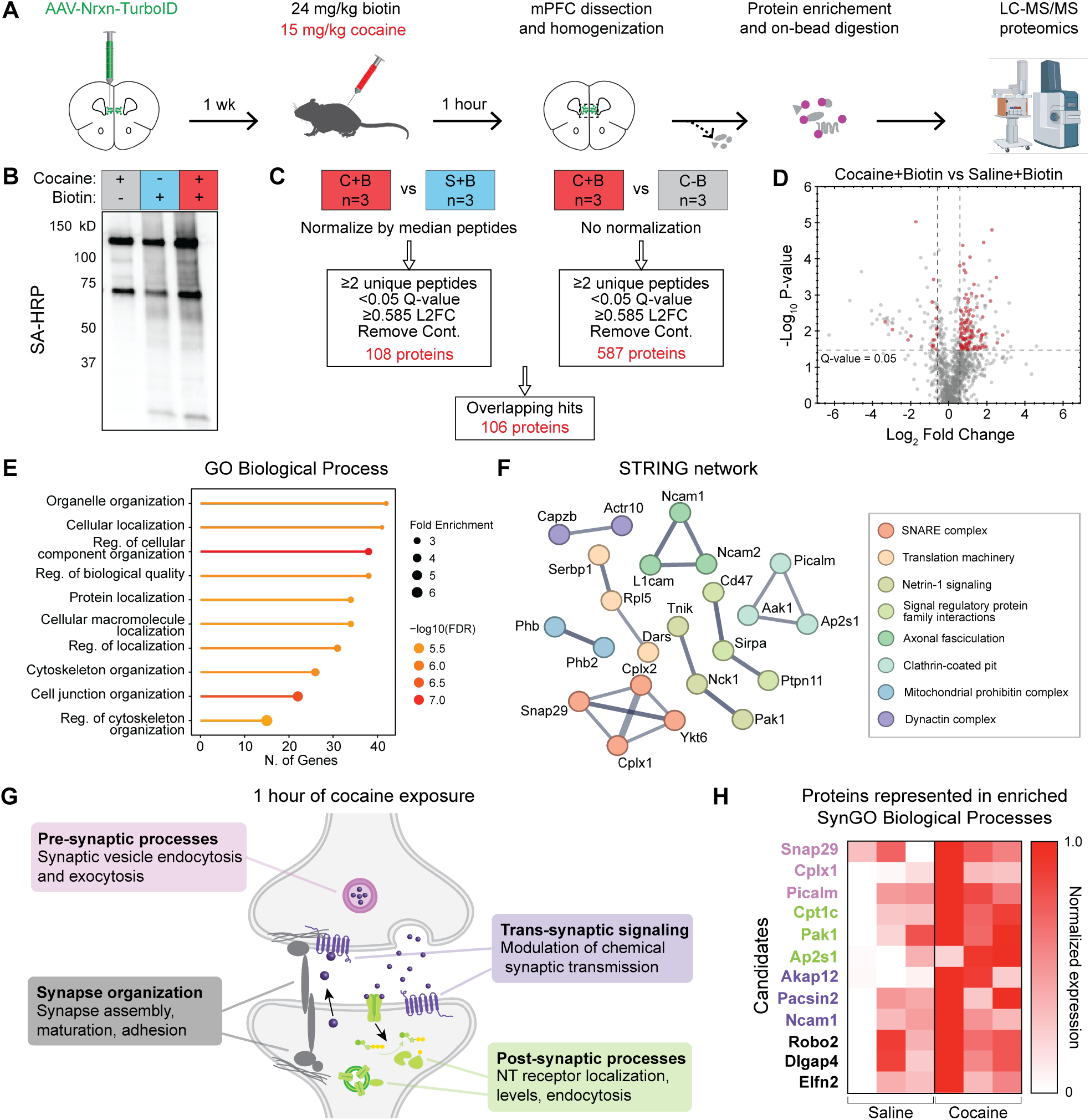
Enrichment of biotinylated proteins in mPFC neurons after acute drug exposure. (**A**) Schematics for TurboID experiments following cocaine injection. AAV2/1-Syn-Nrxn-TurboID was injected bilaterally in mPFC. 1 week later, mice were injected once with either 24 mg/kg biotin or vehicle and 15 mg/kg cocaine or saline. 1 hour later, mice were flushed with PBS through transcardial perfusion, and the brains were collected for mPFC microdissection. Tissue was homogenized and proteins were enriched using magnetic streptavidin beads for LC-MS/MS. Samples were processed and analyzed using DIA proteomics. **(B)** Western blot showing SA-HRP and anti-V5 staining of biotin-enriched samples. **(C)** Schematic for filtering analysis to identify proteins upregulated in response to drug. **(D)** Volcano plot showing results of cocaine+biotin vs saline+biotin differential protein expression analysis from mPFC cell bodies. Vertical dashed lines correspond to L2FC 0.585. **(E)** Top 10 GO enrichment analysis terms for cellular compartment performed using the filtered proteins upregulated after drug in panel (D). **(F)** STRING network analysis of filtered drug-upregulated proteins. **(G)** Schematic depicting enriched synaptic biological function terms associated with hits in panel (D) identified using SynGO. (**H**) Heatmap of min-max normalized raw quantity of exemplar proteins represented in the 4 categories shown in panel (G). See also **Figures S13-S15**.

We further analyzed the drug-upregulated hits using a GO enrichment analysis for biological processes (**Fig. 6E**) and a STRING^43,44^ network analysis (**Fig. 6F, Table S7**). We found that the drug-upregulated hits were involved in processes such as cellular organization and localization, and synaptic signaling. SynGO^45^ enrichment analysis for biological processes specific to synapses showed that the drug-upregulated hits were involved in regulating pre-synaptic vesicle endocytosis and exocytosis; modulation of chemical synaptic transmission; synapse structural assembly, maturation, and adhesion; and post-synaptic localization, regulation, and endocytosis of neurotransmitter receptors (**Fig. 6G, Table S8**). These hits included pre-synaptic proteins such as Snap29, Cplx1, and Picalm; post-synaptic receptor trafficking proteins such as Cpt1c, Pak1, and Ap2s1; trans-synaptic signaling proteins Akap12, Pacsin2, and Ncam1; and synaptic scaffolding proteins like Robo2, Dlgap4, and Elfn2 (**Fig. 6H**).

## DISCUSSION

We developed new membrane-targeted TurboID variants and demonstrated their application for subcellularly targeted neuronal proteomics and drug-gated proteomics in vivo in the mouse brain. Our two tested variants, Nrxn-TurboID and TurboID-CAAX, differ in the mechanism by which they are targeted to the membrane. The truncated Nrxn3b is a larger 115 amino acid protein (13 kDa) that is naturally targeted to the membrane, bringing the fused TurboID with it. CAAX2 is a smaller 22 amino acid motif (2.2 kDa) that is added to the C terminal of TurboID to promote membrane localization through prenylation. Because of this difference in targeting mechanism, we expect that there are differences in the precise subcellular localization of these two constructs in neurons. Indeed, via IHC we could sometimes detect low levels of cytosolic expression of Nrxn-TurboID itself. This could be due to post-translational modifications that take place to the truncated Nrxn3b, such as regulated intramembrane proteolysis^46^, which would result in low levels of cytosolic localization. Note that both variants could result in biotinylation of proteins present within the protein sorting/trafficking pathway, given that newly synthesized Nrxn-TurboID and CAAX-TurboID could be undergoing active sorting and transport during the biotin injection.

We found that TurboID-CAAX resulted in fewer proteins detected overall compared to Nrxn-TurboID. However, the TurboID-CAAX proteome maintained a higher sensitivity for true positive proteins at matched proteome sizes compared to the Nrxn-TurboID proteome, highlighting their complementary strengths for different experimental goals. Thus in certain contexts (when trying to maximize the total protein detection in small compartments such as axons), it may be better to use Nrxn-TurboID; but in other contexts (when trying to maximize membrane specificity), it may be desirable to use TurboID-CAAX. Although a different MS instrument was used to collect the TurboID-CAAX dataset, it is unlikely that this explained the reduction in proteins, given that our Western blot analysis also showed reduced biotinylated protein levels above background in the TurboID-CAAX samples (with levels too low in the mPFC-to-NAc terminals to proceed with proteomics; **Figure S7**). However, future studies would be needed to perform a direct side-by-side comparison of the two different variants.

We also showed that a 1-2 hour labeling window with two IP biotin injections is sufficient for TurboID to tag membrane-targeted proteins above background endogenous biotin levels. This validation suggests that any TurboID experiment performed in the mouse brain, for a diverse set of applications, can be performed with orders of magnitude shorter timescales than is typically used. This importantly allows us to limit the overall exposure of exogenous biotin to the mouse brain and body. While a prior study generating a transgenic TurboID mouse^24^ reported that prolonged biotin exposure did not cause detectable adverse side effects in mice, nor did it alter the baseline electrophysiological properties of neurons in the brain, it is still possible that biotin may cause unintended side effects in cellular proteomes or physiology.

Additionally, we showed that Nrxn-TurboID is sensitive enough to be functional at the terminals of mPFC projections in the NAc; although it may require pooling of biological samples for sparser projections such as to the PAG, as none of these candidates survived the adjusted Q-value thresholding. Nonetheless, we compared our membrane-targeted proteins found in PFC-to-NAc or PFC-to-PAG terminals, and determined several hits that were exclusively present in either brain region. To probe the specificity of these hits for each projection, we compared our proteomics dataset to a previously published RNA-seq dataset^28^ that compared the transcriptome of mPFC neurons projecting to either the NAc or PAG. This comparison likely underestimates the total specificity of our mPFC-to-NAc vs -PAG proteome, given that 1) gene expression does not always correlate to protein expression, and 2) the RNA-seq dataset does not capture locally translated RNAs in axon terminals that could be associated with some of our protein hits that were not detected in the RNA-seq dataset. Nonetheless, the strong correspondence between the datasets from the two different modalities support our approach for projection-specific axonal proteomics in vivo. These data, along with our shorter labeling windows, open future possibilities for studying proteomic changes between various neuronal projections involved in driving a range of different behaviors. Future biological applications using these approaches however will also require antibody-based validation when possible.

While here we used ROC analysis to characterize our membrane-proximal protein hits against known neuronal membrane proteins (true positives) and nuclear proteins (false positives), future studies can develop a nuclear-targeted TurboID AAV construct that can be used in the same studies as our Nrxn-TurboID construct. This would enable a more direct comparison and refinement of “membrane hits” that are present at levels above proteins labeled in a different subcellular neighborhood.

Finally, we showed that Nrxn-TurboID can be used to capture acute drug-induced proteomic changes in mice that were given an acute injection of cocaine and biotin. We showed our labeling approach captured ∼106 proteins engaged at mPFC neuronal cell membranes in response to the drug. Many of these proteins could be mapped onto the SynGO database, highlighting the specificity of our approach for enriching neuronal membrane-related proteins. Furthermore, many of the identified proteins (**Table S8**) were represented by significantly enriched synaptic activity-related biological processes (**Fig.6G, H**). Our study thus serves as a proof of principle that this approach can be used to determine stimulus-induced changes in protein levels at a given subcellular location; however, additional studies are needed to determine whether these proteins are specific to cocaine-induced signaling or instead reflect a more general response to stimulant-driven increases in neuronal activity.

Future studies using our platform can determine comprehensive proteomic changes involved in acute exposure to different drugs or toxins, within genetically and/or anatomically defined cell bodies or axons. Identifying protein networks altered in neuronal circuit pathways implicated in animal models may provide more accurate druggable targets^47^ for treating neuropsychiatric and neurological disorders.

### Limitations of the study

Although here we have demonstrated the compatibility of TurboID with an acute 1-hour labeling window in vivo, peroxidase-based enzymes such as APEX2^6^ are still the fastest option for labeling proximal proteins within minutes (albeit with the requirement of ex vivo labeling). Further improvements to TurboID are thus needed to improve the catalytic efficiency of labeling. We also employed AAVs instead of transgenic TurboID mouse lines, to facilitate the flexible targeting of TurboID to different brain regions; however, a key limitation is that virally mediated overexpression of TurboID requires careful imaging-based and protein-based validation to confirm proper localization of the tools. In addition, differences in virally mediated expression of TurboID across animals may mask more subtle protein level changes across experimental conditions.

Another limitation of the approach presented here is that while the TurboID is fused to the C terminus of the truncated Neurexin-3β membrane targeting motif, we believe that the protein candidates we are reporting represent proteins that are generally present in the same subcellular region surrounding the intracellular membrane, rather than representing true protein-protein interaction partners with the endogenous neurexin protein itself. While in other contexts, TurboID is used at carefully controlled expression levels in cells to identify true protein interaction partners with a given bait protein^18^, at the high, virally-mediated expression levels used here, we believe that TurboID is labeling a broader subset of proteins in close proximity to the membrane, rather than neurexin-interaction partners.

The application of our optimized TurboID variants and shortened labeling protocol provides an opportunity to ask new questions regarding proteomic changes within specific neural circuits that may occur after an exposure to a behavior or drug. While proximity labeling tools can provide vast amounts of new and important hypothesis-generating data, it will be essential to cross-reference and validate the resulting protein lists with orthogonal tools and modalities to investigate biologically relevant questions. Our studies were also performed only in male mice, thus future studies can examine whether there are sex differences in cocaine-induced protein expression, as previously reported^48^.

Finally, here we used only a neuron-specific promoter to drive membrane-targeted expression of TurboID in the mPFC. While this allows specificity of proteins restricted to neuronal membranes, it does not distinguish proteins from different mPFC cell-types^49^. Thus, future studies can use enhancer^50^ or transgenic line^51^ based AAV targeting approaches to identify membrane proteins within specific genetically- and anatomically-defined cell-types.

## RESOURCE AVAILABILITY

### Lead contact

Requests for further information and resources should be directed to and will be fulfilled by the lead contact, Dr. Christina Kim.

### Materials availability

Plasmids generated in this study have been deposited to Addgene (see Key Resources Table for accession numbers).

### Data and code availability

Mass Spectrometry data and search results can be located in the Massive data repository https://massive.ucsd.edu/ using the Dataset number MSV000099974 and proteome exchange https://proteomecentral.proteomexchange.org using the Dataset ID PXD071005. All raw data is publicly available at https://doi.org/10.6084/m9.figshare.30747140. All processed proteomics data is provided in **Tables S1-S5**. Original code is publicly available at https://github.com/tinakimlab/Anguiano2026.

## ACKNOWLEDGMENTS

V5-TurboID-NES_pCDNA3 was a gift from Alice Ting (Addgene plasmid #107169). We thank Himanshu Mali (Princeton University) for assistance with Addgene plasmid preparation. This work was supported by the National Institutes of Health (R21DA059842 to C.K.K., R25NS112130 to M.R., F31MH138072-01A1 to E.L., 1S10OD026918-01A1 to B.S.P., F31DA062491 to E.F., and R35GM119831 to A.S.N.), the Gordon and Betty Moore Foundation (to C.K.K.), the Searle Scholars Program (SSP-2022-107, to C.K.K.), the Arnold and Mabel Beckman Foundation (BYI, to C.K.K.), and the National Science Foundation (Graduate Research Fellowship Program 000895154 to M.A., and Research Traineeship NeuralStorm 2152260 to R.Z.).

## AUTHOR CONTRIBUTIONS

M.A. and C.K.K. conceived the project. R.Z. and M.R. performed molecular engineering, cultured experiments, and analysis. M.A. performed in vivo mouse experiments, proteomics sample preparation, and proteomics analysis. M.R.S. and B.P. performed LC-MS/MS and raw proteomics data analysis. E.L. and J.W. assisted with cocaine experiments. K.P.A. assisted with mouse validation experiments. K.G. assisted with proteomics analysis. C.L. and S.L. generated key viruses for the project. E.F. and A.S.N. performed RNA sequencing dataset reanalysis. M.A., R.Z., and C.K.K. prepared figures and wrote the initial manuscript draft. All authors contributed to reviewing and editing the manuscript. C.K.K. supervised the project.

## SUPPLEMENTAL INFORMATION

**Document S1. Figures S1–S15**

**Table S1 [excel]. Individual sample Log2 quantity protein levels for all experiments**

**Table S2 [excel]. Summary of unfiltered identified proteins for all experiments**

**Table S3 [excel]. Summary of filtered protein candidates for all experiments**

**Table S4 [excel]. Summary of contaminants removed from proteomics experiments**

**Table S5 [excel]. ROC curve analysis**

**Table S6 [excel]. RNA-seq differential expression reanalysis for PFC projection neurons**

**Table S7 [excel]. STRING analysis for Cocaine hits**

**Table S8 [excel]. SynGO biological processes analysis for Cocaine hits**

## STAR★METHODS

### KEY RESOURCES TABLE

The items in the key resources table (KRT) must also be reported alongside the description of their use in the method details section. Literature cited within the KRT must be included in the references list. Please **do not edit the headings or add custom headings or subheadings** to the KRT. We highly recommend using [RRIDs](https://scicrunch.org/resources) as the identifier for antibodies and model organisms in the KRT. To create the KRT, please use the template below or the [KRT webform](https://star-methods.com/). See the more detailed [Word table template](http://www.cell.com/pb-assets/journals/research/cell/methods/table-template1.docx) document for examples of how to list items.

| REAGENT OR RESOURCE | SOURCE | IDENTIFIER |
| --- | --- | --- |
| **Antibodies** |  |  |
| Streptavidin-Alexa647 | Invitrogen | #S32357 |
| Mouse anti-V5 | Invitrogen | #R96025 |
| Donkey anti-mouse Alexa488 | Abcam | #ab150105 |
| Streptavidin-HRP | Invitrogen | s911 |
| Anti-mouse-HRP | Bio-Rad | 170-6516 |
| **Bacterial and virus strains** |  |  |
| AAV2/1-Syn-Nrxn-V5-TurboID-NES (3.52e11 vg/ml) | This paper | N/A |
| AAV2/1-Syn-V5-TurboID-CAAX2 (2.26e10 vg/ml) | This paper | N/A |
| **Chemicals, peptides, and recombinant proteins** |  |  |
| Biotin | Sigma | B4639 |
| Trypsin | Millipore Sigma | T2610 |
| Protease inhibitor cocktail | Sigma-Aldrich | P8849 |
| PMSF | G-Biosciences | 786-055 |
| **Deposited data** |  |  |
| Proteomics dataset | This paper | MassIVE: MSV000099974;<br>ProteomeXchange: PXD071005 |
| PFC, NAc and PAG RNAseq dataset | Lui et al. <sup>28</sup> | NCBI GEO: GSE161936 |
| Experimental models: Cell lines |  |  |
| HEK293T | ATCC | CRL-3216 |
| Rat hippocampal tissue | Brain Bits | SDEHP |
| Experimental models: Organisms/strains |  |  |
| C57/BL6 male mice, Strain #:000664 | Jackson Laboratory | IMSR_JAX:000664 |
| Recombinant DNA |  |  |
| pAAV-Syn-V5-TurboID-NES | This paper | Addgene 250171 |
| pAAV-Syn-Nrxn-V5-TurboID-NES | This paper | Addgene 250172 |
| pAAV-Syn-Nrxn-CIBN-V5-TurboID-NES | This paper | Addgene 250174 |
| pAAV-Syn-CD4-CIBN-V5-TurboID-NES | This paper | Addgene 250173 |
| pAAV-Syn-V5-TurboID-CAAX1 | This paper | Addgene 250175 |
| pAAV-Syn-V5-TurboID-CAAX2 | This paper | Addgene 250176 |
| Software and algorithms |  |  |
| Spectronaut | Biognosys Schlieren | https://biognosys.com/software/spectronaut/ |
| Prism | Graphpad software | https://www.graphpad.com |
| ShinyGO | Ge et al. <sup>34</sup> | https://bioinformatics.sdstate.edu/go/ |
| DeepVenn | Hulsen <sup>52</sup> | https://www.deepvenn.com |
| STRING | Szklarczyk et al. <sup>44</sup> | https://string-db.org/ |
| ImageJ | Schneider et al. <sup>53</sup> | https://imagej.net/ij/index.html |
| SynGO | Koopmans et al. <sup>45</sup> | https://www.syngoportal.org |
| Visual Studio Code | Microsoft | https://visualstudio.microsoft.com/ |
| IPython Notebook | IPython | https://ipython.org/ |
| MATLAB R2025a | Mathworks software | https://www.mathworks.com/products/matlab.html |
| Claude Code | Anthropic | https://code.claude.com/ |
| Other |  |  |
| Pierce Streptavidin magnetic beads | Thermo Scientific | PI88817 |

### METHOD DETAILS

#### Cloning

All constructs developed in this study are listed in Key Resources Table. Vector backbones were double digested with restriction enzymes (New England BioLabs; NEB) following standard digestion protocols. Q5 polymerase (M0494S, NEB) was used to amplify PCR fragments. Backbone vectors and PCR inserts were purified using gel electrophoresis followed by gel extraction (28706, Qiagen) and were assembled using Gibson reactions (E2611S, NEB). Plasmids were heat shock transformed into NEB Stable Competent Escherichia coli (C3040H, NEB) following the manufacturer’s protocol. Plasmids were amplified using a Plasmid Miniprep Kit (27106, Qiagen). Constructs were cloned from the following source plasmids or sequences: V5-TurboID-NES (gift from Alice Ting, Addgene plasmid #107169); CD4-sTb(C)-M13-GFP (Addgene plasmid #219784); Nrxn3b-CIBN-Nav1.6-MKII-hLOV1-TEVcs(ENLYFQ/M)-tTA-VP16 (gift from Alice Ting, Addgene plasmid #163029).

#### Mammalian cell culture and crude AAV2/1 production

Human Embryonic Kidney 293T cell (CRL-3216, ATCC) were cultured as a monolayer in Dulbecco’s Modified Eagle Medium (DMEM; D5796-500ML, Sigma-Aldrich), supplemented with 10% Fetal Bovine Serum (FBS; F1051-500ML, Sigma-Aldrich) and 1% (v/v) penicillin/streptomycin (P/S; 15070063, 5,000 U/mL, Life Technologies) (complete DMEM). Cells were cultured in 100 mm TC treated dishes (353003, Falcon) at 37°C and 5% CO_2_ and subcultured when they reached 80%–90% confluence using Trypsin (T2610-100ML, Sigma-Aldrich).

HEK293T cells at ∼80% confluency in 100 mm dishes were transfected with 2.3 µg AAV vector, 1.92 µg AAV1 plasmid, 1.92 µg AAV2 plasmid, 4.6 µg DF6 AAV helper plasmid, and 69 µL PEI Max solution per construct and were incubated for 24 hr. The cell culture media was completely replaced with 10 ml fresh media and the cells were incubated for another 48 hr for virus production. After the incubation, the cell culture conditioned media (supernatant) from the 100 mm dishes was collected, filtered through 0.45 µm pore size filter (9914-2504, Cytiva) and aliquoted for cultured neuron infection.

#### Primary neuronal culture, AAV infection, and biotin treatment

E18 rat hippocampal neurons were dissociated, plated, and infected following the manufacturer’s protocols (SDEHP, BrainBits). Neurons were plated on treated 35 mm glass bottom dishes (D35-14-1-N, Cellvis) and maintained at 37°C, 5% CO_2_. The culture media was replaced 24 hr after the plating with complete Neurobasal Plus medium supplemented with 0.01% (v/v) Gentamicin, 0.75% (v/v) Glutamax, 2% (v/v) B27 Plus to stop glial growth. Subsequently, ∼50% of the media was replaced every 3-4 days. At DIV6, neurons were infected with crude supernatant AAVs of different variants of membrane-TurboID variants by replacing half of the culture media. The dishes were incubated for another ∼two weeks in the incubator prior to experiments. Prior to stimulation, a portion of the culture media from each dish was taken out and saved in microcentrifuge tubes. The biotinylation mixtures were generated by adding the biotin solution to the saved culture media to the desired concentration. The biotinylation mixtures were then added back to each dish and the neurons were incubated at 37°C for 10 min for membrane-TurboID labeling. For +biotin conditions, we used a final concentration of 50 µM biotin. Neurons were then fixed and stained as described below.

#### Immunohistochemistry in cultured neurons

After incubation for the indicated time, the solution was removed in each dish. Neurons were washed once with warm complete DMEM and twice with warm DPBS (D8537-500ML, Sigma-Aldrich). Neurons were subsequently fixed with 4% (v/v) paraformaldehyde (PFA; sc-281692, Santa Cruz Biotechnology) in DPBS at room temperature for 10 min followed by two washes with DPBS. After that, neurons were permeabilized with ice-cold methanol at - 20°C for 8 min followed by two washes with DPBS at room temperature. Neurons were incubated with blocking solution (1% w/v BSA; BP1600-100, Fisher Scientific, in DPBS) for 45 min at room temperature; followed by a 1 hr incubation in primary antibody solution at room temperature (1:1,000 mouse anti-v5; R96025, Invitrogen). Neurons were washed twice with DPBS and incubated with secondary antibody (1:1,000 donkey anti-mouse 568; ab175472, Abcam) and streptavidin-Alexa Fluor647 (1:5,000; S32357, Invitrogen) for 30 min at room temperature. Neurons were washed twice with DPBS and imaged by fluorescence microscopy.

#### Fluorescence imaging and analysis in cultured neurons

Fluorescence images were taken with Keyence BZ-X810 fluorescence microscope with an 80W metal halide lamp as the fluorescent light source and a PlanApo 10X air objective lens (NA 0.45) or PlanApo 40X air objective lens (NA 0.95). Expression of membrane-TurboID was visualized by Alexa Fluor568 using 545/25 nm excitation filter and 605/70 nm emission filter. Biotinylated proteins were labeled by SA-647 and visualized by 620/60 nm excitation filter and 700/75 nm emission filter. A CCD camera was used to acquire fluorescence images. Images were analyzed by custom scripts in ImageJ and MATLAB. Where indicated, confocal imaging was performed using a Zeiss LSM 800 confocal microscope (Carl Zeiss) equipped with 561 and 640 nm lasers, and a 100X (NA 1.3) oil immersion objective. 35 mm glass bottom dishes with a 14 mm micro-well and #1 cover glass (0.13mm-0.16mm) were used for high resolution confocal imaging. Fluorescence images were collected with 1024 x 1024-pixel resolution and with a pixel dwell time of 0.52 µsec/pixel. Images were acquired using a PMT detector and emission filter ranges of 560–640 nm and 645–700 nm for Alexa Fluor568 and Alexa Fluor647 detection, respectively, for best signal. All images were collected and processed using ZEN software (Carl Zeiss). For membrane-TurboID fluorescence characterization, we quantified the mean fluorescence intensity of each FOV. Typically, 9-10 FOVs were imaged for a given experimental condition. We calculated the mean SA-647/V5 cell ratios for each FOV. This analysis can normalize for any difference in expression levels of the tool across cells and conditions.

#### Wildtype mice

All animal-related procedures were approved by the UC Davis Institutional Animal Care and Use Committee. Wildtype C57BL/6J male mice 5-7 weeks old were obtained from Jackson laboratory (Strain #000664). The mice used in this study were on a 12 h light/12 h dark cycle with ad libitum access to food and water. Animals were housed in the vivarium under standard conditions for mice (temperature 72°F, humidity range 40–50%).

#### AAV2/1 preparation for infection in vivo

For purification and concentration of AAV2/1s used for in vivo expression, HEK293T cells were cultured to ∼80% confluency in three T150 flasks for each construct. Each flask was transfected with 5.2 µg of the expression vector, 4.35 µg of AAV1 plasmid, 4.35 µg of AAV2 plasmid, 10.4 µg of DF6 AAV helper plasmid, and 130 µL of PEI solution. After transfection, cells were incubated for 48 hours and washed with cold PBS prior to collection. For virus purification and concentration, the HEK293T cells were pelleted at 800 × g for 10 minutes, then resuspended in 20 mL of 100 mM TBS (100 mM NaCl, 20 mM Tris, pH 8.0). A 10% sodium deoxycholate solution (Sigma) was added to a final concentration of 0.5%, along with benzonase nuclease (Sigma) at 50 units/mL. The suspension was incubated at 37 °C for 1 hour, then clarified by centrifugation at 3,000 × g for 15 minutes. The supernatant was applied to a HiTrap heparin column (Cytiva) pre-equilibrated with 10 mL TBS using a peristaltic pump. After viral binding, the column was washed with 10 mL TBS via the pump, followed by 1 mL of 200 mM TBS and 1 mL of 300 mM TBS using a syringe. Viral particles were eluted sequentially with 1.5 mL of 400 mM TBS, 3.0 mL of 450 mM TBS, and 1.5 mL of 500 mM TBS. The eluate was concentrated using a 15 mL centrifugal filter unit (100 K MWCO, Amicon) spun at 2,000 × g for 2 minutes, reducing the volume to approximately 500 µL. The virus was further concentrated using a 0.5 mL centrifugal unit (100 K MWCO, Amicon) to a final volume of 100–200 µL. Viral titers were quantified using the AAVpro Titration Kit Ver.2 (Takara Bio Inc). For titration, 2 µL of AAV sample was mixed with 1 µL DNase I (Takara), 2 µL 10× DNase I buffer (Takara), and 15 µL dH_2_O, and incubated for 15 minutes at 37 °C. DNase I was then inactivated by heating at 95 °C for 10 minutes. Subsequently, 20 µL of lysis buffer (Takara) was added and incubated at 70 °C for 10 minutes. The resulting AAV solution was diluted 50-fold, and 5 µL was used as the PCR template together with 0.5 µL primer mix (50×, containing 5 µL forward primer, 5 µL reverse primer, and 15 µL dH_2_O), 7 µL dH_2_O, and 12.5 µL 2× TB Green Premix Ex Taq II (Takara) for real-time PCR. A standard curve was generated using serial dilutions of the positive control solution (Takara) for absolute quantification. Viral titers were determined based on the standard curve.

#### Stereotaxic surgeries and in vivo biotin labeling in mice

Briefly, 5-7 week-old mice were maintained under anesthesia with 2% isoflurane and immobilized on an RWD stereotaxic apparatus. For immunohistochemistry experiments, unilateral intracranial injections in mPFC (ML +/-0.5, AP +1.98, DV -2.25) were performed with 800 nL of AAV2/1-Syn-Nrxn-V5-TurboID-NES or AAV2/1-Syn-V5-TurboID-CAAX2 for experimental and control groups. Injections were performed with a Hamilton syringe and WPI syringe pump, at a rate of 150 nL/min. For proteomics experiments, bilateral intracranial injections in the mPFC were performed with 500 nL of AAV2/1-Syn-Nrxn-V5-TurboID-NES or AAV2/1-Syn-V5-TurboID-CAAX2 for experimental and control groups. Once the surgery was completed, the incision was closed with tissue adhesive (GLUture), and each mouse received EthiqaXR for analgesia. This AAV2/1 serotype has been previously demonstrated to express broadly in neurons both in cortical and subcortical brain regions (cortex^29,54^, nucleus accumbens^55^, hippocampus^56^, ventral tegmental area^55^).

After a week of viral expression, mice were given one to two 24 mg/kg biotin IP injections from a 10 mM biotin stock solution (Sigma, B4639 dissolved in 10% DMSO). Mice were transcardially perfused one hour after the last biotin injection. Control mice were injected with vehicle solution made of DMSO and DPBS.

#### Immunohistochemistry and fluorescence imaging in mouse brain tissue

For IHC analysis, after biotin labeling, mice were perfused with ice-cold PBS and 4% PFA, and brains were collected and stored in 4% PFA overnight at 4°C. The next day, brains were switched out from PFA and stored in PBS until slicing. 60µm coronal slices were collected from the mPFC, NAc and PAG and placed in wells with PBS and stored in 4°C. Slices were washed in PBS-T for 2 min (3x). Then blocked in 5% Normal Donkey Serum and 0.3% Triton-X (PBS-T) for 1 hour at room temperature. Slices were stained in primary antibody (1:500 dilution, mouse anti-V5, Invitrogen #R96025) in 5% NDS/PBS-T overnight. The next day, slices were washed with PBS-T for 5 minutes (3x) at room temperature. Slices were then stained with secondary antibody (1:1000 dilution, donkey anti-mouse Alexa488, Abcam #ab150105) and streptavidin-Alexa647 (1:1000 dilution, Invitrogen #S32357) in 5% NDS/PBS-T for 1.5 hours at room temperature. Slices were washed with PBS-T for 5 minutes at room temperature and then mounted with DAPI-Fluoromount-G (SouthernBiotech) to adhere coverslips. Images were taken on Keyence BZ-X810 fluorescence microscope (acquisition software v1.1.2) fluorescence microscope.

Confocal imaging was performed using a Carl Zeiss LSM 800 confocal microscope equipped with 488-, 561- and 640-nm lasers, and a ×63 oil immersion objective (NA 1.4). All images were collected and processed using ZEN software v2.3 (Carl Zeiss).

#### Protein extraction

After biotin labeling, mice were perfused with PBS and brains were collected. Using a brain block and pairs of single blades, the mPFC, NAc and PAG were microdissected bilaterally from ∼1-2 mm thick coronal sections. Each brain region was placed in a 1.5 mL tube containing 200 uL of RIPA buffer: 50 mM Tris/HCL, 150 mM NaCl, 1mM EDTA supplemented with protease inhibitor cocktail (Sigma-Aldrich, P8849) and PMSF (G-Biosciences, 786-055). Each sample was homogenized using disposable pestles driven by a mechanical motor. Then, 200 µL of additional Ripa buffer (50 mM Tris/HCL, 150 mM NaCl, 1mM EDTA supplemented with protease inhibitor cocktail and PMSF) was added, followed by 500 µL of RIPA lysis buffer (50 mM Tris/HCL, 150 mM NaCl, 1mM EDTA, 0.4% SDS, 2% Triton x-100, 2% sodium deoxycholate supplemented with protease inhibitor cocktail and PMSF). Each sample was sonicated using 20% Amplitude with 5-second pulses for 30 seconds. Samples were then centrifuged at 13,000 g for 30 min at 4°C. The supernatants were transferred to new tubes, supplemented with 1% SDS, and incubated at 45°C for 45 minutes. Samples were then cooled on ice, and 3% of the total volume was set aside as “whole-cell-lysate” by flash freezing and storing in -20°C. The rest of the sample was incubated with 65 µL Pierce Streptavidin magnetic beads (Thermo Scientific, PI88817) at 4°C overnight with rotation.

#### Biotinylated protein enrichment

Samples were placed on a magnetic rack, and the cleared supernatant was saved as “flow-through” in a 1.5 mL tube and kept on ice. The beads were then washed in 75 µL of 50 mM ammonium bicarbonate for 20 minutes and placed on rotation during each incubation at 4°C (4x). 5% of the volume after the fourth wash was saved in a 1.5 mL tube containing 1 mL of RIPA lysis buffer for analysis using Streptavidin blot. The rest of the sample was prepared for proteomics by adding 2 uL of trypsin (Millipore Sigma, T2610) to each sample and incubated overnight at room temperature on rotation.

#### Protein visualization using Western blot

Following overnight rotation with magnetic beads, 5% of the sample that was saved in a 1.5 mL tube containing 1 mL of RIPA lysis buffer, placed on a magnetic rack, and the clear supernatant was discarded. 30 µL of 3x protein loading buffer supplemented with 20 mM DTT (Fisher, BP172-5) and 2 mM biotin was added to the dry beads. Additionally, 2 µL of 6x Laemmli protein loading buffer (ThermoScientific, J60660.AC) was added to 10 µL of flow-through and 4 uL of 6x protein loading buffer was added to 20 µL of whole cell lysate. All samples were boiled at 95°C for 10 min. Samples were placed back on the magnetic rack and supernatant was collected and stored in a new 1.5 mL tube. Proteins from supernatant, flow-through and whole cell lysate were separated in 10% precast gels (Biorad, #4561035) for 45 minutes at 215 V and transferred to nitrocellulose membrane (Cytiva BioTrace, 66485) for 1 hour at 350 mAmps. Blots were stained with 0.2% Ponceau for 10 min and imaged. Blots were then thoroughly washed with 1x TBST (0.1% Tween-20) and then blocked with 5% nonfat dry milk (bioKEMIX, M0841). To detect V5, blots were incubated in 3% BSA/TBST containing mouse anti-v5 primary (1:10000; Invitrogen, R96025) for 1 hour at room temperature. Membrane was washed three times with TBST for 5 minutes and incubated in 3% BSA/TBST containing anti-mouse-HRP secondary antibody (1:10,000; Bio-Rad 170-6516) for 30 minutes at room temperature. To detect biotinylated proteins, blots were incubated in 3% BSA/TBST containing streptavidin-HRP (1:5000, Invitrogen, s911) for 30 minutes at room temperature. Membranes were washed with TBST and then developed using ECL western blotting substrate (BioRad, 1705062) for 5 min. The membrane blots were imaged on a Bio-Rad Gel Doc XR gel imager. The Western blot images were visualized using ImageJ.

#### On bead digestion

After overnight incubation with trypsin (Millipore Sigma, T2610), samples were placed on a magnetic rack, and the cleared supernatant was collected and placed in a new 1.5 mL tube. The remaining beads were washed again with 50 mM ammonium bicarbonate and placed on a shaker for 20 min at room temperature. The samples containing the beads were placed back in the magnetic rack, and the cleared supernatant was then removed and combined with the first extract. Samples were sent to the UC Davis Proteomic core for LC-MS/MS.

#### LC-MS/MS and Protein identification and quantification

##### For Nrxn-TurboID datasets

For LC, peptides were resolved on a Thermo Scientific Dionex UltiMate 3000 RSLC system using a PepSep analytical column (PepSep, Denmark): 150um x 8cm C18 column with 1.5 μm particle size (100 Å pores), preceded by a Pepmap Neo C18 5UM guard column, and heated to 40 °C. Separation was performed in a total run time of 30 min with a flow rate of 500 μL/min with mobile phases A: water/0.1% formic acid and B: 80%ACN/0.1% formic acid. For MS DIA, peptides were directly eluted onto an Orbitrap Exploris 480 instrument (Thermo Fisher Scientific, Bremen, Germany). Spray voltage was set to 1.8 kV, funnel RF level at 45, and heated capillary temperature at 275 °C. Experiment’s full MS resolution was set to 60,000 at m/z 200 and full MS AGC target was 300% with an IT of 45msec. Mass range was set to 350–1200. AGC target value for fragment spectra was set at 1000%. Mass Windows of 15Da were used with an overlap of 1 Da. Resolution was set to 30,000 and IT to Auto. Normalized collision energy was set at 30%. All data were acquired in profile mode using positive polarity and peptide match was set to off, and isotope exclusion was on. The Thermo Fisher LC/MS system is supported by NIH grant 1S10OD026918-01A1. Initial DIA data was analyzed using Spectronaut 16.2 and 17 (Biognosys Schlieren, Switzerland) using the directDIA workflow with the default settings. Briefly trypsin/P Specific was set for the enzyme allowing two missed cleavages. Fixed Modifications were set for Carbamidomethyl, and variable modification were set to Acetyl (Protein N-term) and Oxidation. For DIA search identification, PSM and Protein Group FDR was set at 0.01%. A minimum of 2 peptides per protein group were required for quantification. Peptide identification was performed by searching against the UniProt *Mus* musculus reference proteome (UP000000589_10090 Mouse 1 entry per gene, 21,986 entries), and a Universal Contaminant Protein FASTA database.

##### For TurboID-CAAX datasets

MS was performed on a hybrid trapped ion mobility spectrometry-quadrupole time of flight mass spectrometer (timsTOF HT, (Bruker Daltonics, Bremen, Germany) with a modified nano-electrospray ion source (CaptiveSpray, Bruker Daltonics). In the experiments described here, the mass spectrometer was operated in PASEF mode. Desolvated ions entered the vacuum region through the glass capillary and deflected into the TIMS tunnel which is electrically separated into two parts (dual TIMS). Here, the first region is operated as an ion accumulation trap that primarily stores all ions entering the mass spectrometer, while the second part performs trapped ion mobility analysis. For DIA parallel accumulation-serial fragmentation (PASEF), the dual TIMS analyzer was operated at a fixed duty cycle close to 100% using equal accumulation and ramp times of 85 ms each. Data-independent analysis (DIA) scheme consisted of one MS scan followed by MSMS scans taken with 36 precursor windows at width of 25Th per 1.09 sec cycle, over the mass range 300-1200 Dalton. The TIMS scans layer the doubly and triply charged peptides over an ion mobility -1/k0-range of 0.7-1.3 V*sec/cm2. The collision energy was ramped linearly as a function of the mobility from 59 eV at 1/K0=1.4 to 20 eV at 1/K0=0.6. Initial DIA data was analyzed using Spectronaut 19.5 (Biognosys Schlieren, Switzerland) using the direct DIA workflow. Trypsin/P Specific was set for the enzyme allowing two missed cleavages. Fixed Modifications were set for Carbamidomethyl, and variable modifications were set to Acetyl (Protein N-term), and Oxidation. For DIA search identification, PSM and Protein Group FDR was set at 0.01%. A minimum of 2 peptides per protein group were required for quantification. Peptide identification was performed by searching against the UniProt *Mus* musculus reference proteome (UP000000589_10090 Mouse 1 entry per gene, 21,986 entries), and a Universal Contaminant Protein FASTA database.

### Secondary proteomics analysis

DIA data were analyzed for differential expression across conditions with Spectronaut 20.4.260109 (Biognosys Schlieren, Switzerland). Run wise imputation strategy was used for all datasets in this paper. See raw data included with the referenced Figshare Dataset for excel sheets of peptides that were imputed. The initial list of candidates (≥ 2 unique peptides, L2FC ≥ 0.585, Q/P-value < 0.05) differed in the following way with and without run-wise imputation:

- mPFC Nrxn-TurboID: 1,918 candidates with run-wise imputation; 1,778 candidates without
- mPFC TurboID-CAAX: 515 candidates with; 177 without
- mPFC-to-NAc terminals Nrxn-TurboID: 1,962 candidates with, 1,511 without
- mPFC-to-PAG terminals Nrxn-TurboID: 191 exploratory candidates with, 155 without
- mPFC cocaine+biotin vs cocaine-biotin Nrxn-TurboID: 613 candidates with, 524 without
- mPFC cocaine+biotin vs saline+biotin Nrxn-TurboID: 109 candidates with, 71 without

Cross-run normalization was turned off for all datasets comparing biotin and no biotin controls. Cross-run normalization was turned on (global normalization using median levels of only peptides identified across 100% of runs) for the cocaine+biotin versus saline+biotin dataset. All identified proteins are listed in **Table S1** (individual runs) and in **Table S2** (averaged by condition). Proteins from each data set were identified as candidates if they had two or more unique peptides, a fold-change ≥ 1.5 (Log2 fold-change ≥ 0.585) and a Q-value less than 0.05. For the PFC-to-PAG Nrxn dataset only, a P-value threshold of <0.05 was used (**Table S3**). Common contaminants found in proteomics data were identified and excluded^57^ (**Table S4**).

To map the mPFC Nrxn-TurboID candidates to the 10X Isocortex Allen Mouse Brain Cell Atlas^49^, the Seurat^58^ AddModuleScore was used, to create a “proteomics score” for each cell in the Allen Brain Atlas, based on how highly it expressed the set of genes corresponding to the PFC Nrxn-TurboID candidates. The score was calculated as score_cell = mean(query_gene_expression) – mean(expression-matched background gene expression).

To calculate the ROC curves, sensitivity, and specificity metrics for the proteomes, a true positive list was used for proteins identified in mouse striatal neurons using LCK-APEX2^13^ (lymphocyte-specific protein tyrosine kinase) for Nrxn-TurboID and TurboID-CAAX mPFC proteomes. For PFC-to-NAc and PFC-to-PAG Nrxn-TurboID terminal proteomes, a true positive list was curated from SynGO^45^ using presynaptic membrane associated terms. A false positive list was curated from the same LCK-APEX2 study, from a nuclear H2B-APEX2 dataset, along with additional nuclear terms curated from UniProtKB. All terms used for ROC analysis are in **Table S5**.

#### Proteomics data visualization

Heatmaps of normalized expression of top candidates comparing experimental and control groups were created in Graphpad Prism v10.5.0. Proteins from each data set were identified as candidates if they had two or more unique peptides, a L2FC ≥ 0.585 and a Q-value < 0.05. All upregulated candidates were run through ShinyGO 0.85.1 for gene ontology analysis with FDR cut off 0.05 and all other settings were kept as default. All upregulated candidates in our cocaine vs saline dataset were run through STRING v12.0 to perform a protein-protein interaction network analysis (see **Table S7**). The network was generated using the physical subnetwork. Interactions supported by text mining, experimental data and curated database annotations were included. A high-confidence interaction score threshold (≥0.700) was applied. Analysis was limited to interactions among the queried proteins. To identify functional modules within the network, Markov Cluster Algorithm (MCL) clustering was performed using an inflation parameter of 3. SynGO was used to determine biological GO enrichment terms associated with cocaine hits, requiring a minimum of 3 genes per term (see **Table S8**).

#### Comparison to single-cell RNA sequencing dataset for axonal projections

Data was downloaded from NCBI GEO GSE161936 from ref ^28^. The “Fig3Rbp4retro” object was used and the metadata tag “Cre.line” distinguished projections from PFC. To obtain gene counts, the RNA counts from each neuron were 1) normalized then 2) the mean of the normalized counts was taken for each gene across all cells within a projection (e.g. Cav_NAc or Cav_PAG). The normalization occurred within each single cell in which the raw number of reads for a gene was divided by the total number of reads of all genes in that cell, multiplied by a scaling factor of 10,000, then natural-log transformed. A differential expression analysis was run between cells identified as projecting from PFC-to-NAc and cells projecting from PFC-to-PAG. A Wilcoxon test was performed using the Seurat^58^ “FindMarkers” function, including only genes that expressed in a minimum of 5% of cells in either group and with on average at least a 0.05-fold difference (log2-scale) between the two groups of cells. These parameters were selected to be in line with the defaults and to account for cell numbers and the high sequencing depth that this study was able to achieve. P-values were subject to a strict Bonferroni correction using all the genes in the dataset. Data is in **Table S6**.

### QUANTIFICATION AND STATISTICAL ANALYSIS

All statistical analysis details can be found in legends of figures. Statistical analyses including Mann-Whitney *U* test and two-way ANOVA were performed in GraphPad Prism v9.0 (GraphPad Software). Significance was defined as *p<0.05, **p<0.01, ***p<0.001, ****p<0.0001 for the defined statistical test (NS, P ≥ 0.05). Analysis scripts and codebase generation in iPython Notebooks were assisted by Claude Code extension (v2.1.2; Anthropic, 2026) for Visual Studio Code v1.122.1.

