## Supplemental Figures for "Acute in vivo proximity labeling for membrane targeted proteomics in neuronal circuits"

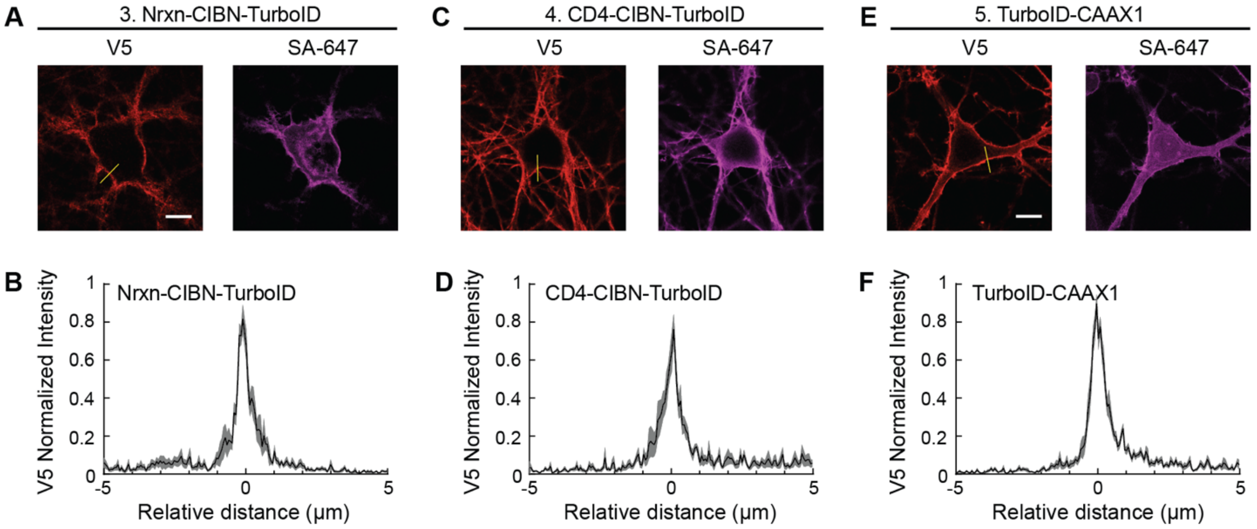


**Figure S1. Confocal imaging of additional membrane-TurboID variants**. **(A)** Example 100x images of anti-V5 and SA-647 staining of cultured neurons expressing Nrxn-CIBN-TurboID and treated with 50 μM biotin for 10 minutes, then fixed and stained for anti-V5 (on TurboID) and SA-647. The yellow line indicates pixels quantified to quantify TurboID membrane localization for summary data in panel B. Scale bar, 10 μm. **(B)** Quantification of the mean anti-V5 membrane localization across neurons (*n* = 10 FOV, data plotted as mean +/- SEM). **(C-F)** Same as panels (A-B), except for cultured neurons infected with CD4-CIBN-TurboID (**C-D**) or TurboID-CAAX1 (**E-F**).

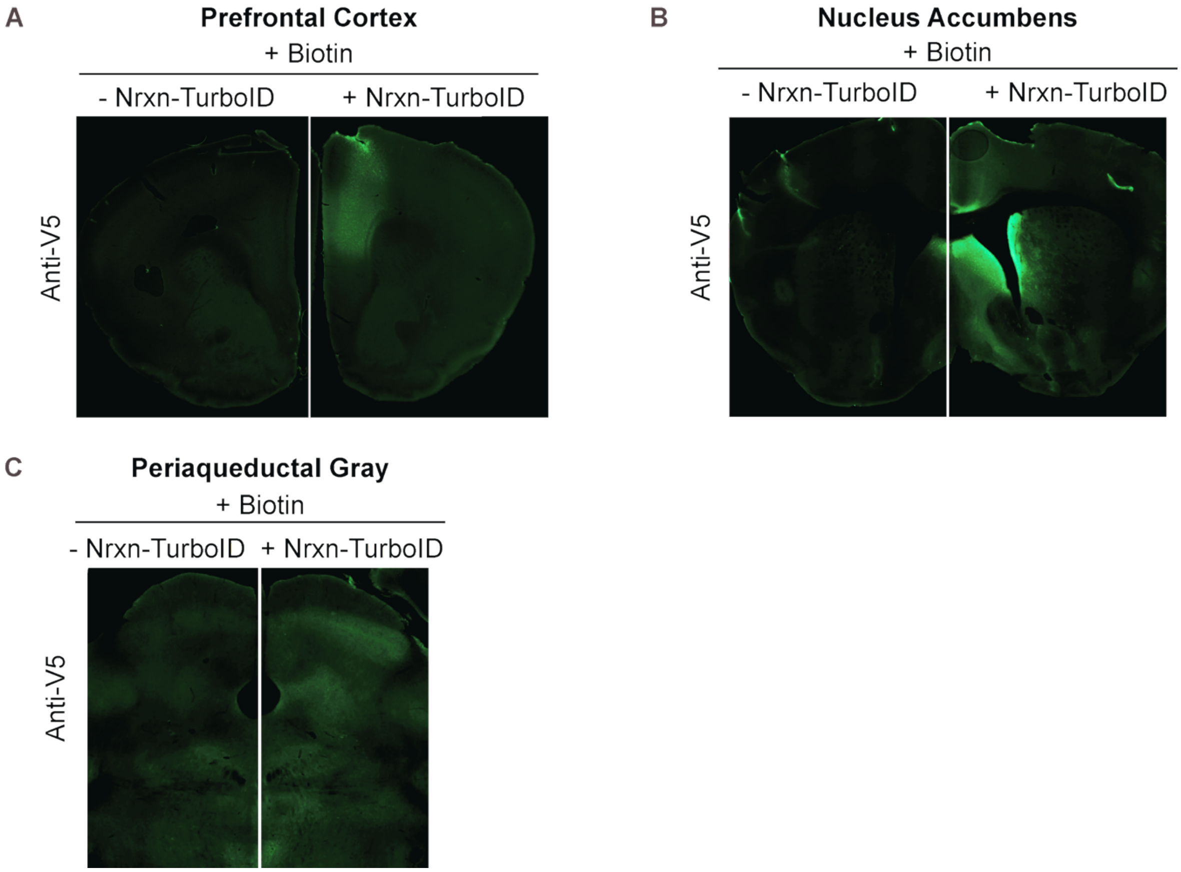


**Figure S2. Representative images of viral targeting in mouse tissue**. **(A)** Entire coronal section showing AAV2/1-Syn-Nrxn-TurboID-V5 staining after injection of 800 nL in the mPFC. The uninjected left hemisphere and injected right hemisphere from the same mouse is shown. **(B-C)** Coronal sections showing expression of AAV2/1-Syn-Nrxn-TurboID-V5 staining in mPFC axons projecting to the nucleus accumbens or periaqueductal gray in the right hemisphere.

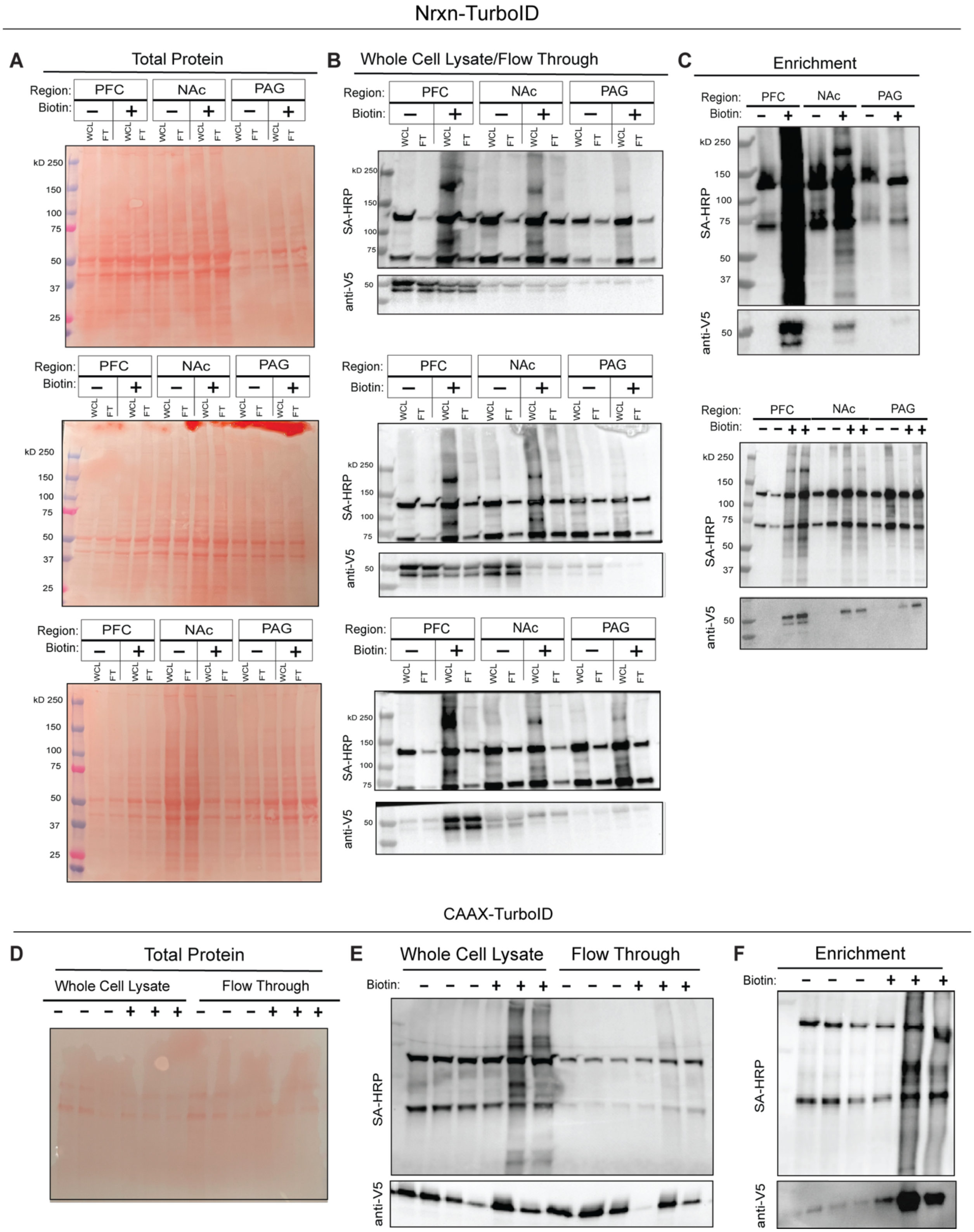


**Figure S3. Full Western blots for protein detection across all mPFC replicates**. **(A)** mPFC, mPFC-to-NAc, and mPFC-to-PAG Nrxn-TurboID samples were run on a 10% gel and transferred onto nitrocellulose membrane. A 0.2% Ponceau stain was used to identify total protein for each replicate following flow-through (FT) and whole cell lysate (WCL) collection. **(B)** Membrane blot containing WCL and FT for mPFC, NAc and PAG samples were stained with SA-HRP for visual confirmation of biotinylated proteins and anti-V5 for visual confirmation of tool expression for each replicate. **(C)** Enrichment was performed using magnetic streptavidin beads for each replicate. Post enrichment, blot was stained for SA-HRP and anti-V5. **(D)** mPFC TurboID-CAAX samples were run on a 10% gel and transferred onto nitrocellulose membrane. A 0.2% Ponceau stain was used to identify total protein for each replicate following FT and WCL collection. **(E)** Membrane blot containing WCL and FT for mPFC samples were stained with SA-HRP and anti-V5. **(F)** Post enrichment, blot was stained for SA-HRP and anti-V5.

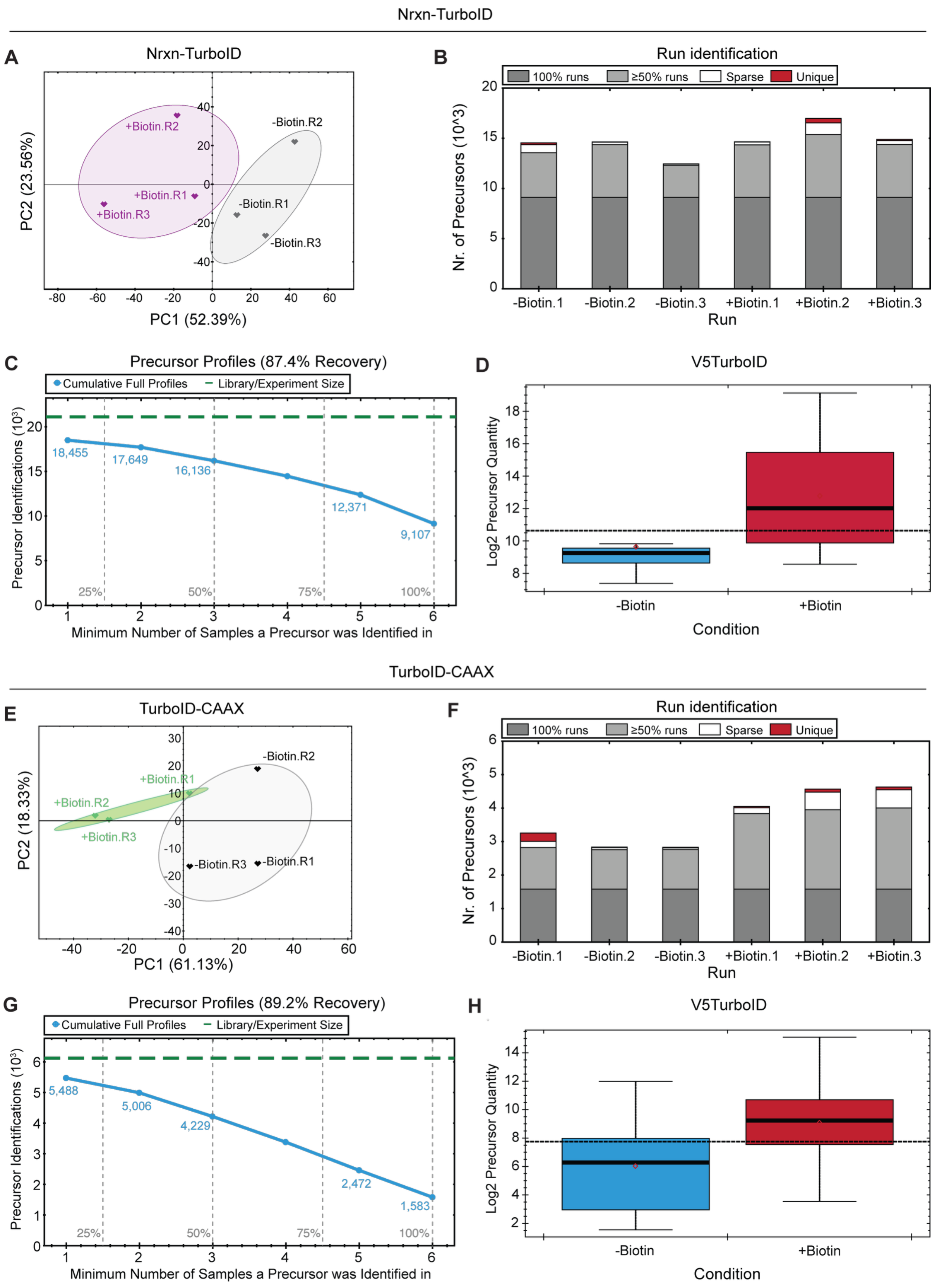
Nrxn-TurboID
A
Nrxn-TurboID
+Biotin.R2
-Biotin.R2
+Biotin.R1
+Biotin.R3
-Biotin.R1
-Biotin.R3
PC2 (23.56%)
PC1 (52.39%)
B
Run identification
100% runs
≥50% runs
Sparse
Unique
Nr. of Precursors (10^3)
-Biotin.1
-Biotin.2
-Biotin.3
+Biotin.1
+Biotin.2
+Biotin.3
Run
C
Precursor Profiles (87.4% Recovery)
Cumulative Full Profiles
Library/Experiment Size
18,455
17,649
16,136
12,371
9,107
25%
50%
75%
100%
Precursor Identifications (10³)
Minimum Number of Samples a Precursor was Identified in
D
V5TurboID
Log2 Precursor Quantity
-Biotin
+Biotin
Condition
TurboID-CAAX
E
TurboID-CAAX
-Biotin.R2
+Biotin.R1
+Biotin.R2
+Biotin.R3
-Biotin.R3
-Biotin.R1
PC2 (18.33%)
PC1 (61.13%)
F
Run identification
100% runs
≥50% runs
Sparse
Unique
Nr. of Precursors (10^3)
-Biotin.1
-Biotin.2
-Biotin.3
+Biotin.1
+Biotin.2
+Biotin.3
Run
G
Precursor Profiles (89.2% Recovery)
Cumulative Full Profiles
Library/Experiment Size
5,488
5,006
4,229
2,472
1,583
25%
50%
75%
100%
Precursor Identifications (10³)
Minimum Number of Samples a Precursor was Identified in
H
V5TurboID
Log2 Precursor Quantity
-Biotin
+Biotin
Condition

**Figure S4. Additional metrics for mPFC TurboID proteomics samples. (A)** Principal component analysis of mPFC Nrxn-TurboID proteomics data shows separated clustering between experimental samples (+biotin, purple) and control samples (-biotin, grey). **(B)** Number of precursors identified in 100% of runs (dark gray), in at least 50% of runs (light gray), in more than 1 but less than 50% of runs (white), or only in a single run (red). **(C)** Cumulative number of precursors identified per minimum number of runs a precursor was identified in. **(D)** Precursor quantity specifically searching for the V5-TurboID in control and experimental samples. **(E-H)** Same as panels (A-D), except for mPFC samples from mice injected with AAV2/1-Syn-TurboID-CAAX.

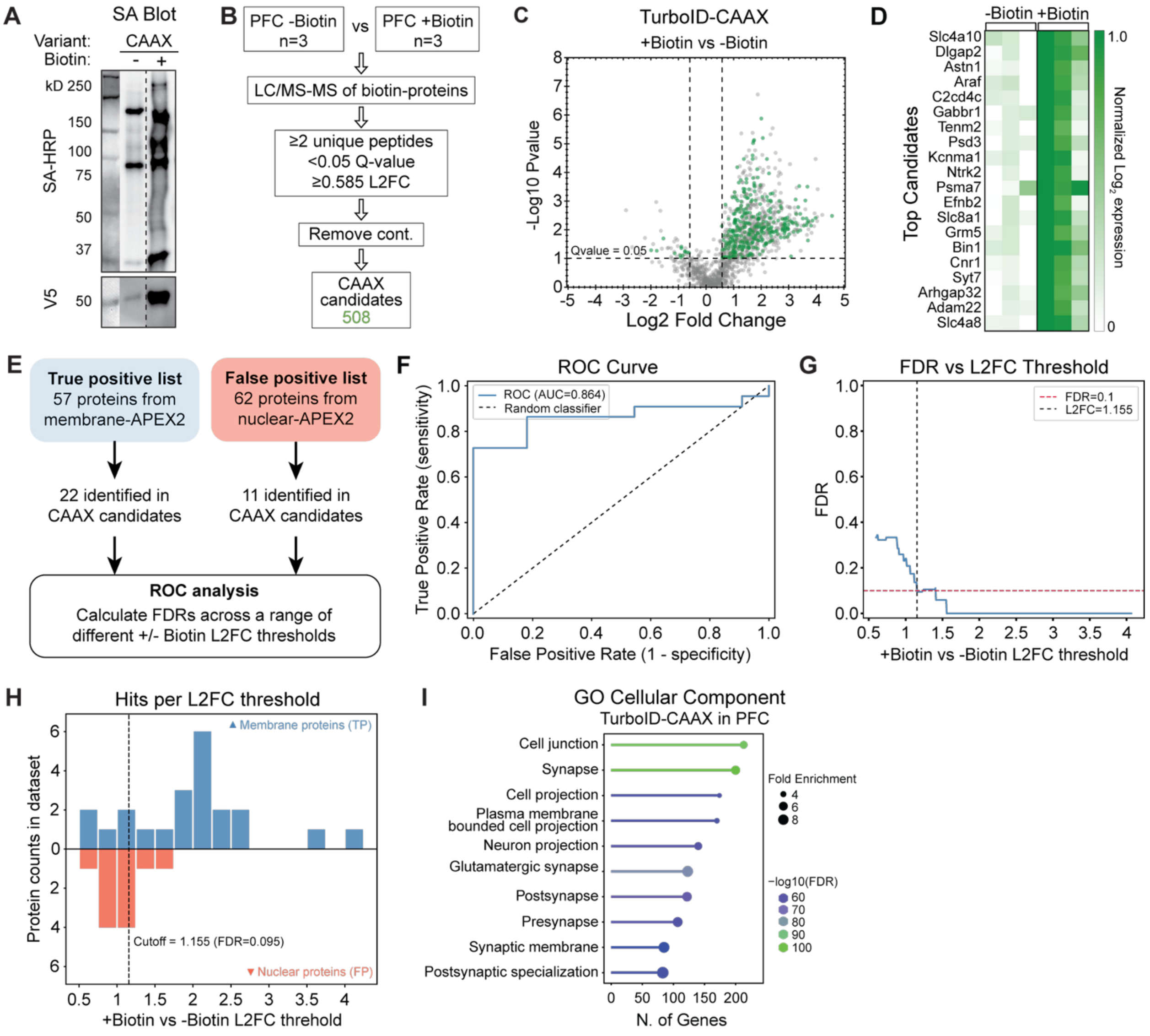


**Figure S5. In vivo protein labeling and proteomics using TurboID-CAAX in mPFC neurons**
**A)** Mice were injected with AAV2/1-Syn-TurboID-CAAX bilaterally, and samples were treated exactly as those from mice injected with AAV2/1-Syn-Nrxn-TurboID in Figure 3. Western blot showing representative SA-HRP and anti-V5 staining of biotin-enriched samples. Dashed vertical line means non-continuous lanes on the same blot are shown. See **Figure S4** for full uncut blots. **(B)** Workflow for data collection and filtering steps. We performed three biological replicates per experimental condition. Proteins were filtered and identified as candidates if they were detected with ≥ 2 unique peptides and had a +/-Biotin L2FC ≥ 0.585 with a Q-value < 0.05. **(C)** Volcano plot showing results of +/- biotin differential protein expression analysis for TurboID-CAAX. **(D)** Heatmap of min-max normalized $\log_2$ expression data for the top 20 candidates with the largest +/-Biotin fold change in the CAAX-TurboID mPFC dataset. **(E)** Schematic depicting strategy for ROC analysis. The same TP and FP lists as in Figure 4 were used. **(F)** ROC curve showing TP rate and RP rate determined using different L2FC thresholds. **(G)** Plot of the FDR versus +/-Biotin L2FC threshold, with the dashed lines indicating the L2FC at which the FDR<0.1. **(H)** Histogram showing the number of TP (blue) and FP (red) proteins present at different +/-Biotin L2FC thresholds. **(I)** Top 10 enriched GO Cellular Compartments determined from the 387 candidates above the L2FC cutoff indicated in panel (H). See also **Table S5.**

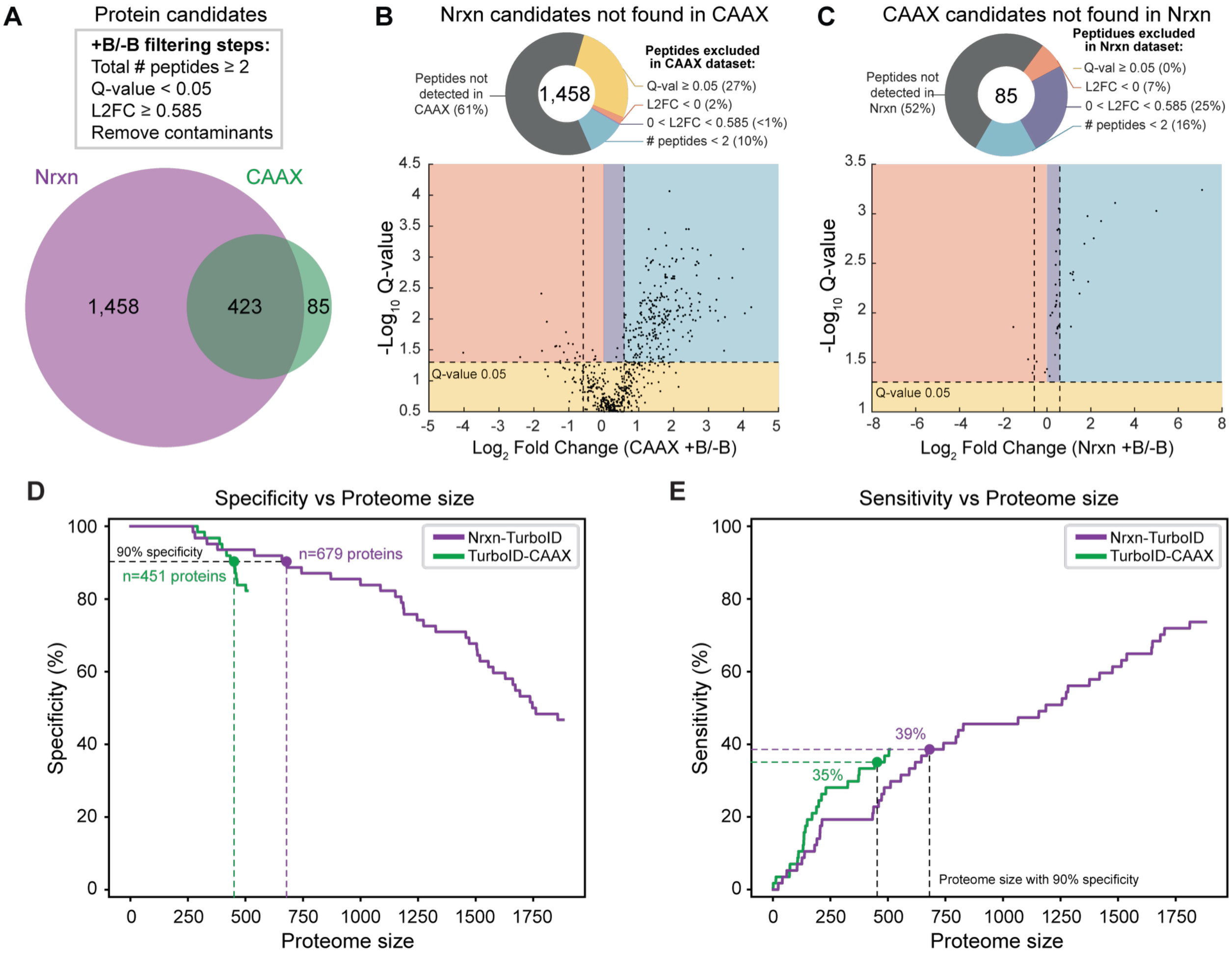


**Figure S6. Comparison of proteomes from mPFC Nrxn-TurboID and TurboID-CAAX datasets. (A)** Diagram showing overlapping and exclusive filtered candidates between Nrxn-TurboID and TurboID-CAAX. **(B)** Analysis of “Nrxn-only” candidates in the entire unfiltered CAAX dataset. Candidates were either not detected in the CAAX LC-MS/MS runs (grey; not plotted); were detected but were not significantly up-regulated between the +/- biotin conditions (yellow); significantly down-regulated in the +/-Biotin condition (orange); significantly up-regulated but below the 1.5-fold enrichment threshold (purple); or significantly up-regulated above threshold, but detected with only 1 unique peptide (blue). **(C)** same as panel B, except analysis of “CAAX-only” candidates in the entire unfiltered Nrxn dataset. **(D)** Analysis of specificity versus increasing proteome size (proteins sorted by +/-Biotin L2FC) for both the Nrxn and CAAX datasets (using the FP nuclear proteome list from Figure 4). The proteome sizes at 90% specificity are shown with dashed vertical lines. **(E)** Analysis of sensitivity versus increasing proteome size (proteins sorted by +/-Biotin L2FC) for both the Nrxn and CAAX datasets (using the TP LCK membrane proteome list from Figure 4). The sensitivities at the proteome sizes corresponding to 90% specificity (from panel (D)) are shown with dashed horizontal lines.

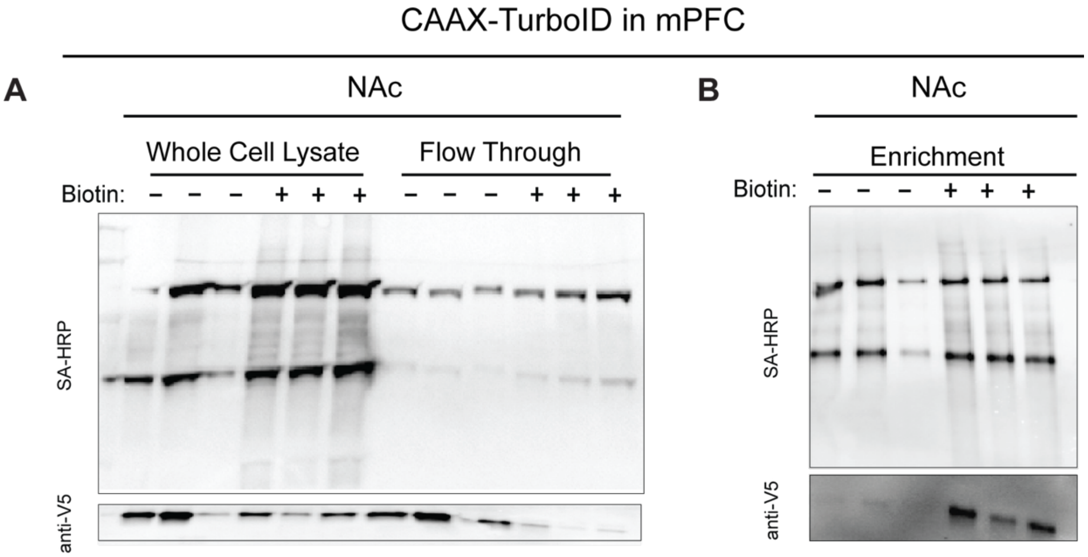


**Figure S7. Western blots showing low sensitivity for CAAX-TurboID in mPFC terminals**. **(A)** The NAc of mice expressing TurboID-CAAX in mPFC were microdissected out and after tissue was lysed, samples were run on a 10% gel and transferred onto nitrocellulose membrane. Blot containing WCL and FT for NAc samples were stained with SA-HRP for visual confirmation of biotinylated proteins and anti-V5 for visual confirmation of tool expression. **(B)** Enrichment was performed using magnetic streptavidin beads for each replicate for NAc samples. Post enrichment, blot was stained for SA-HRP and anti-V5. Given the low levels of biotinylated proteins detected in the mPFC-to-NAc TurboID-CAAX samples, we chose not to proceed with proteomics for the axonal terminal samples.

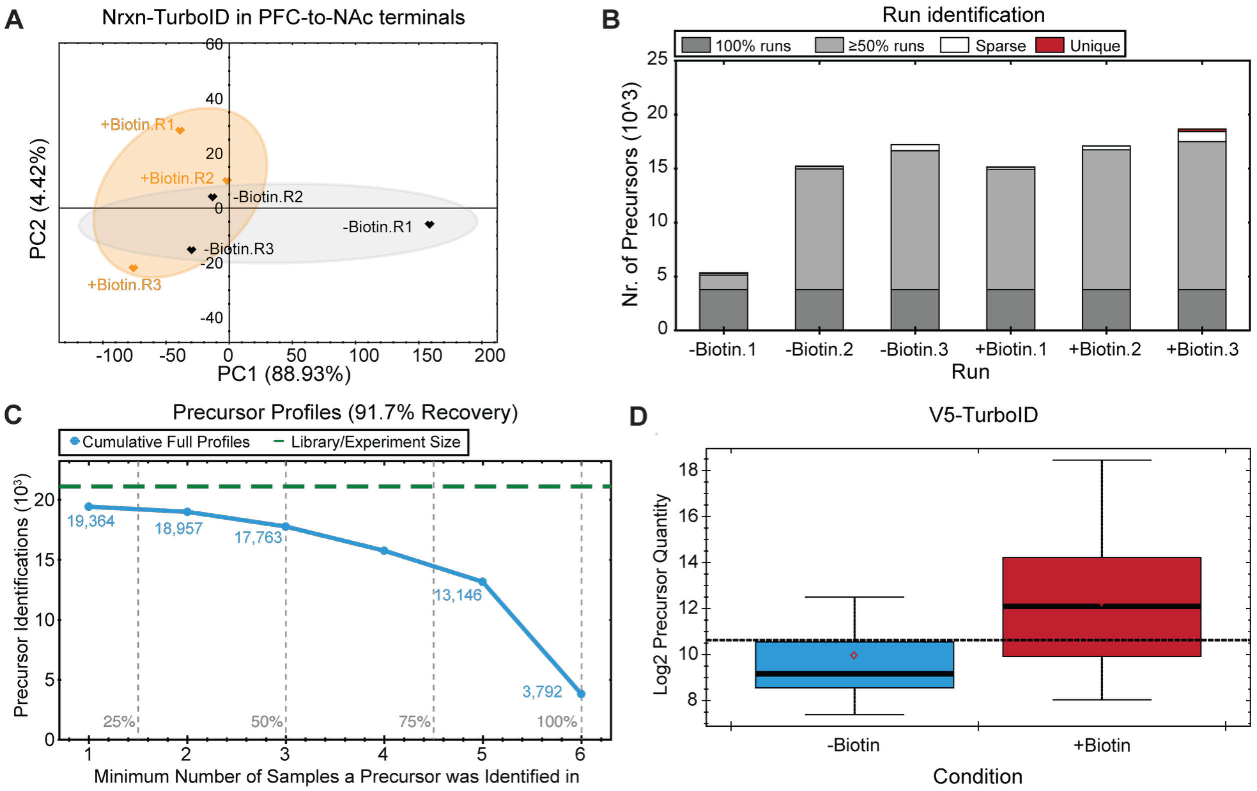


**Figure S8. Additional metrics for mPFC-to-NAc terminals with Nrxn-TurboID proteomics samples. (A)** Principal component analysis of mPFC-to-NAc terminals with Nrxn-TurboID proteomics data shows separated clustering between experimental samples (+biotin, orange) and control samples (-biotin, grey). **(B)** Number of precursors identified in 100% of runs (dark gray), in at least 50% of runs (light gray), in more than 1 but less than 50% of runs (white), or only in a single run (red). **(C)** Cumulative number of precursors identified per minimum number of runs a precursor was identified in. **(D)** Precursor quantity specifically searching for the V5-TurboID in control and experimental samples.

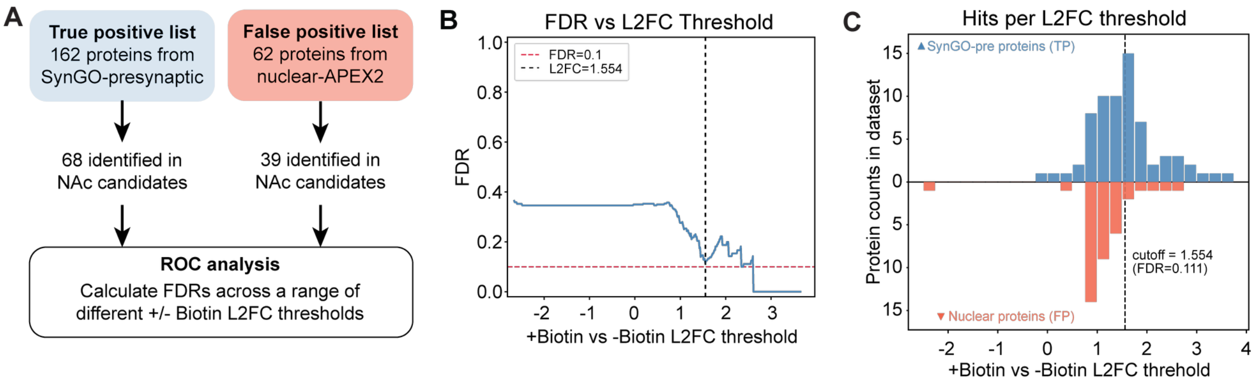


**Figure S9. ROC analysis of mPFC-to-NAc terminals with Nrxn-TurboID. (A)** Schematic depicting strategy for ROC analysis of mPFC-to-NAc terminals Nrxn-TurboID data shown in Figure 5. TP list was taken from SynGO presynaptic terms, and the same FP list was used as in Figure 4. **(B)** Plot of the FDR versus +/-Biotin L2FC threshold, with the dashed vertical line indicating the L2FC at which the FDR=0.111. **(C)** Histogram showing the number of TP (blue) and FP (red) proteins present at different +/-Biotin L2FC thresholds. See also **Table S5.**

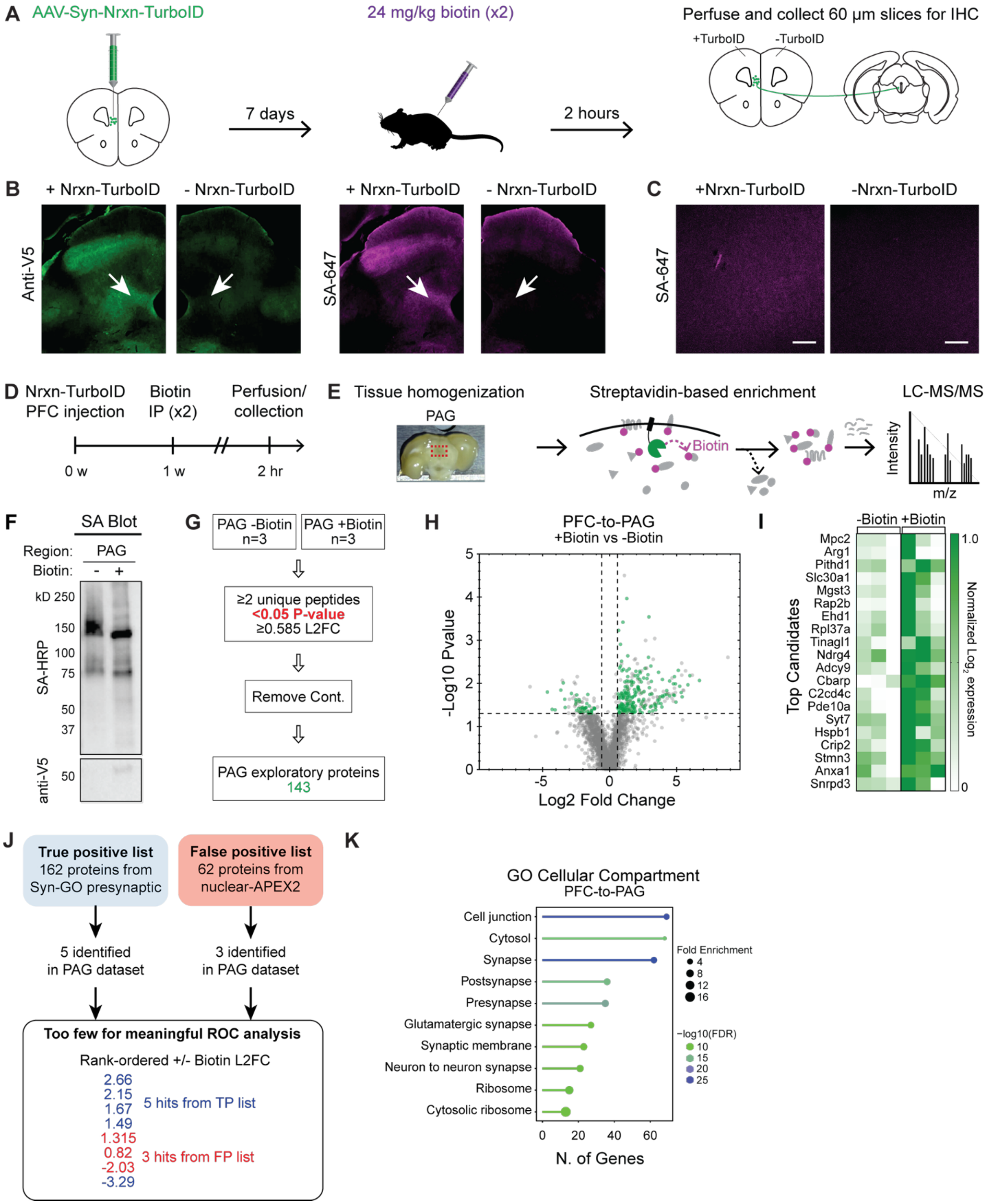


**Figure S10. Exploratory in vivo protein labeling and proteomics of mPFC-to-PAG neuron terminals using Nrxn-TurboID.** ***(A)*** Schematic depicting experimental protocol. Mice were injected unilaterally in the mPFC with AAV2/1-Syn-Nrxn-V5-TurboID. After 1 week, they were injected with two IP biotin (24 mg/kg) or vehicle injections, one hour apart. The brains were then collected for IHC. **(B)** 10x images of the injected and uninjected hemispheres from same mouse, showing the PAG with anti-v5 and SA-647 staining. White arrows point to regions microdissected for proteomics. **(C)** Confocal images of SA-647 staining in PAG. Scale bar, 20 μM. **(D-E)** Schematics for TurboID experiments. AAV2/1-Syn-Nrxn-TurboID was bilaterally injected into mPFC. 1 week later, mice were given two IP biotin (24 mg/kg) or saline injections, one hour apart. Mice were flushed with PBS through transcranial perfusion, and then brains were collected for PAG microdissection. Protein extraction was performed, followed by streptavidin magnetic bead enrichment. After on-bead digestion, peptides were analyzed using DIA proteomics. **(F)** Western blot showing SA-HRP and anti-V5 staining of biotin-enriched samples. **(G)** Workflow for data collection and filtering steps. Note that here, no proteins survived adjusted Q-value <0.05 thresholding, so only unadjusted P-value <0.05 exploratory candidates are shown. **(H)** Volcano plot showing results of +/-Biotin differential protein expression analysis. **(I)** Heatmap of min-max normalized $\log_2$ expression data for the top 20 candidates with the largest +/-Biotin fold change. **(J)** Schematic depicting strategy for ROC analysis. The same TP and FP lists as in Figure 5 were used. However, too few candidates were identified among the PAG dataset to perform a meaningful ROC curve analysis. **(K)** Top 10 GO enrichment analysis terms for cellular compartment performed using the filtered 143 exploratory protein candidates in PAG.

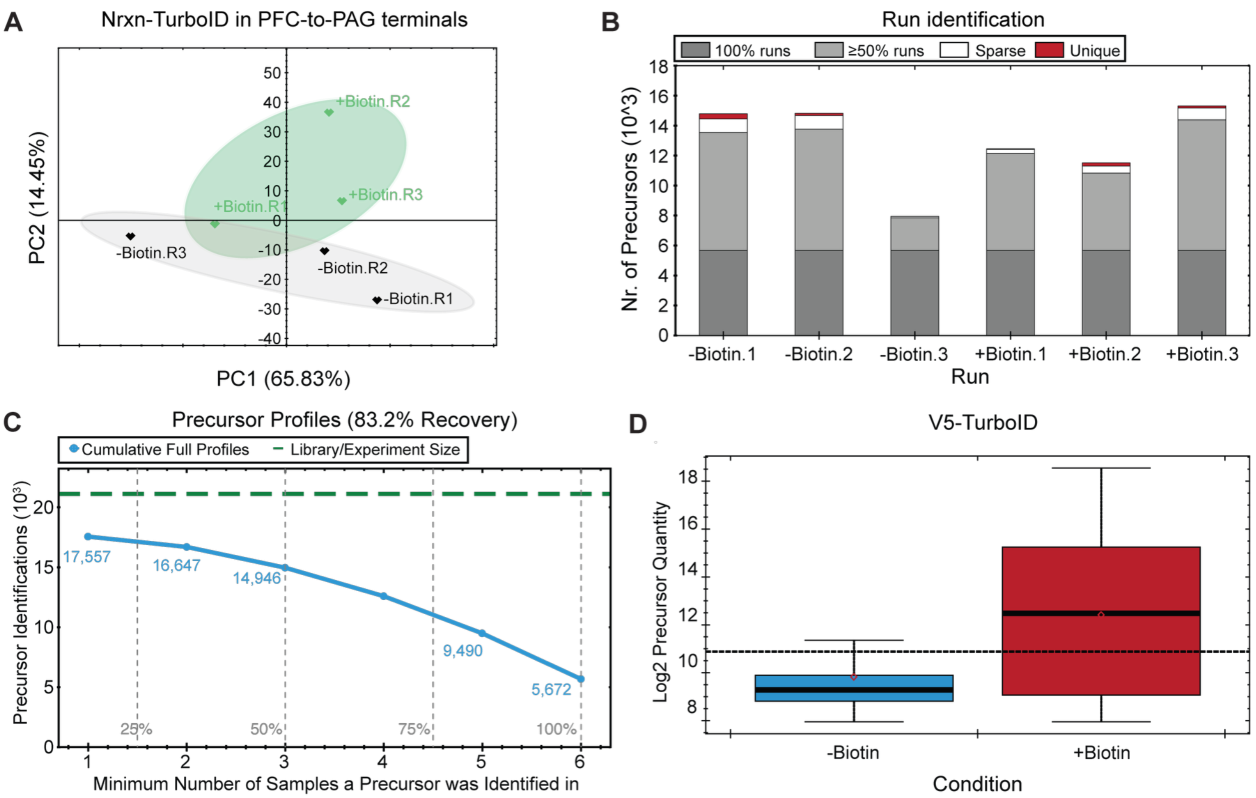


**Figure S11. Additional metrics for mPFC-to-PAG terminals with Nrxn-TurboID proteomics samples. (A)** Principal component analysis of mPFC-to-PAG terminals with Nrxn-TurboID proteomics data shows separated clustering between experimental samples (+biotin, green) and control samples (-biotin, grey). **(B)** Number of precursors identified in 100% of runs (dark gray), in at least 50% of runs (light gray), in more than 1 but less than 50% of runs (white), or only in a single run (red). **(C)** Cumulative number of precursors identified per minimum number of runs a precursor was identified in. **(D)** Precursor quantity specifically searching for the V5-TurboID in control and experimental samples.

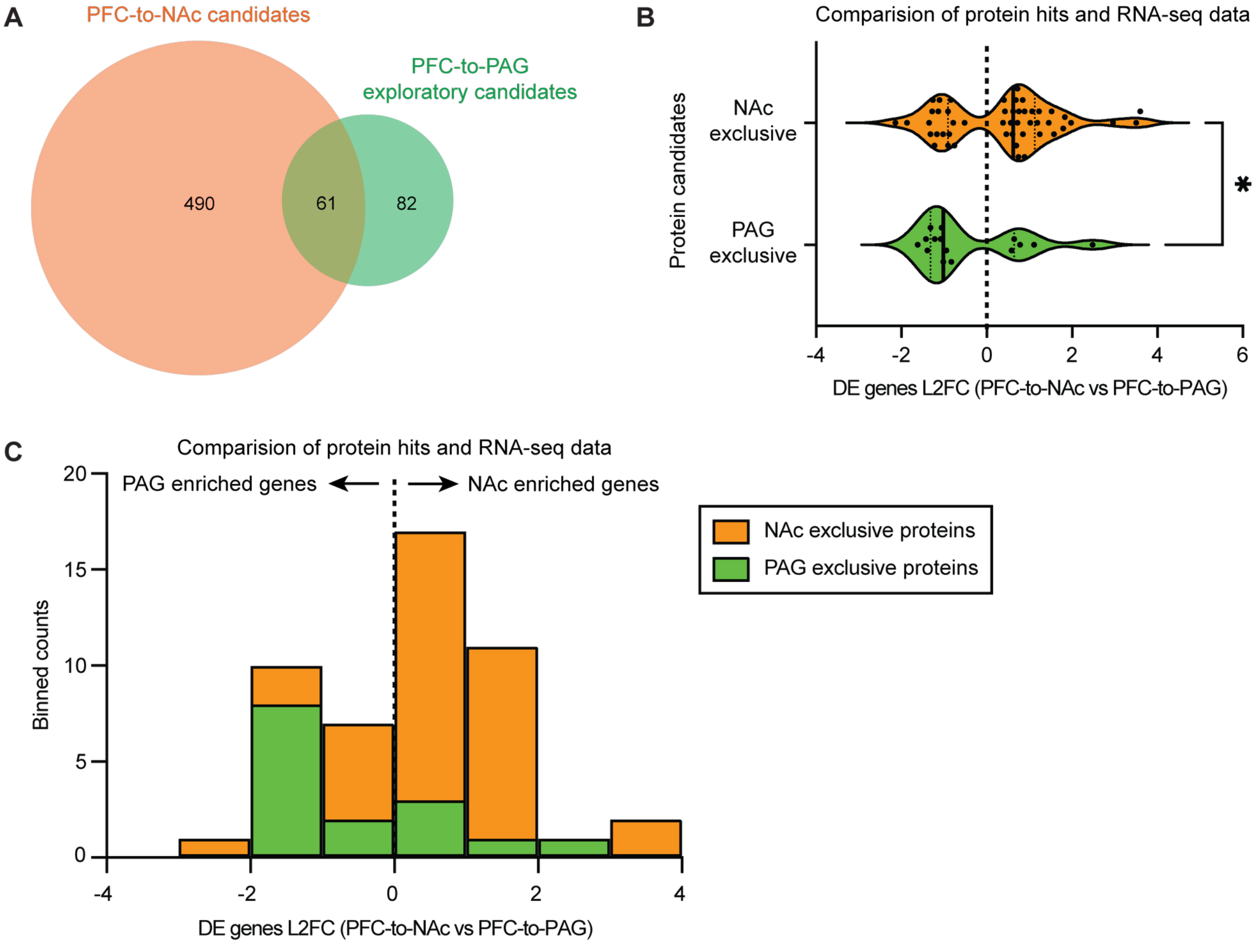


**Figure S12. Comparison of mPFC-to-NAc and mPFC-to-PAG candidates. (A)** Diagram showing overlapping and exclusive filtered candidates between PFC-to-NAc and PFC-to-PAG datasets. **(B)** Volcano plots showing the L2FC RNA expression of DE genes between mPFC-to-NAc and mPFC-to-PAG neurons from a published single cell sequencing dataset. Only DE genes that corresponded to NAc exclusive or PAG exclusive proteins are plotted. **(C)** Histogram counts of the data shown in panel B. Mann-Whitney U test, $^*p<0.05$. See also **Table S6**.

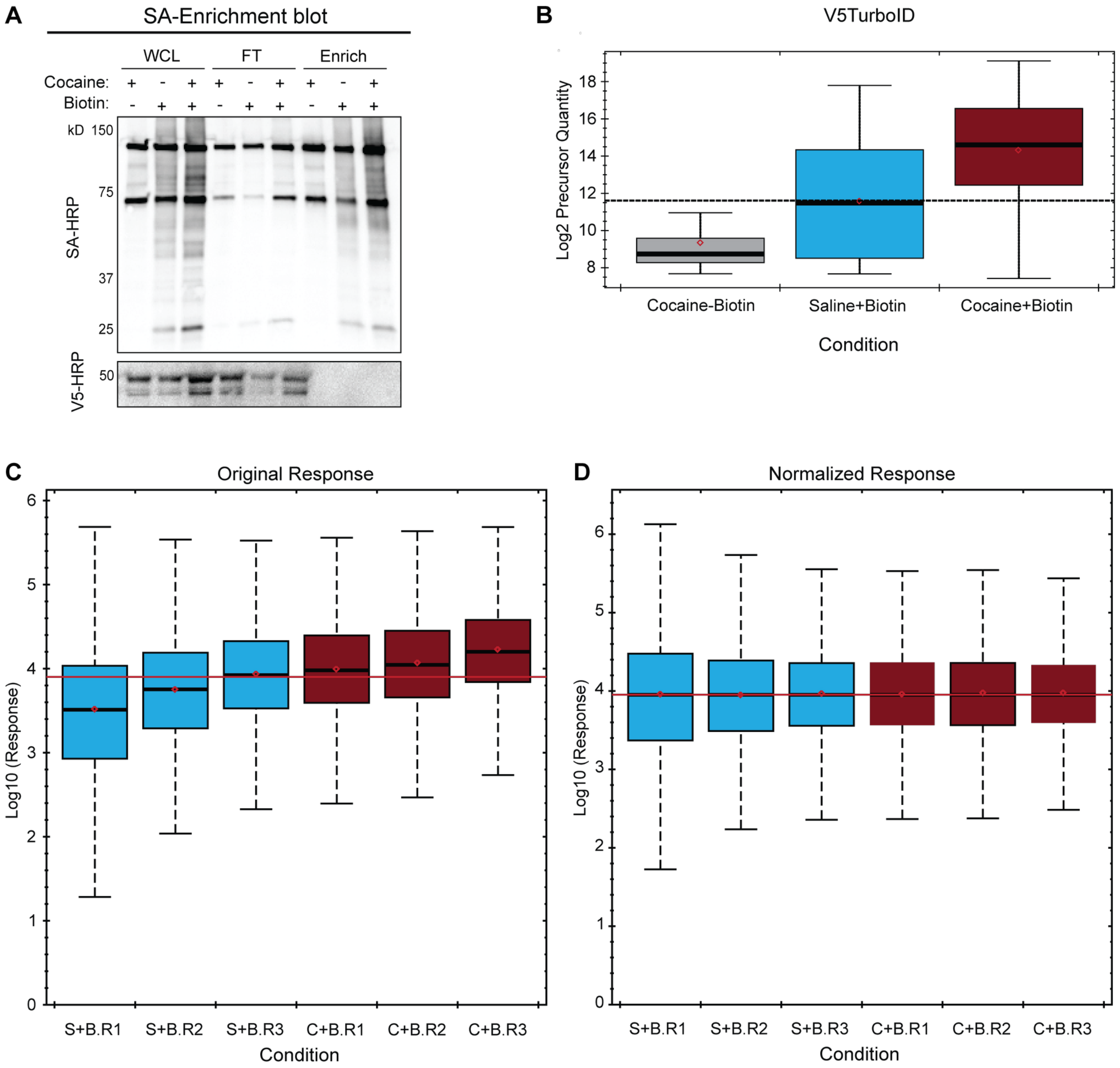


**Figure S13. Western blot and normalization of samples for acute cocaine proteomics experiment. (A)** Samples were run on a 10% gel and transferred onto nitrocellulose membrane. Blot containing WCL, FT and Enrichment samples were stained with SA-HRP for visual confirmation of biotinylated proteins. Note that given the relatively small amount of sample we ran on the gel, we did not observe strong V5 labeling in the enriched samples (although enrichment could be detected at the peptide level using LC-MS/MS). We also only ran 1 sample on a Western blot, in an effort to conserve samples for maximal LC-MS/MS detection. **(B)** Precursor quantity specifically searching for the V5-TurboID in control and experimental samples. **(C)** Quantity of total peptides identified across each Saline+Biotin and Cocaine+Biotin sample prior to normalization. **(D)** Quantity of normalized total peptide levels across each Saline+Biotin and Cocaine+Biotin sample after normalizing according to the global median peptide level in each run, using only peptides detected in 100% of all runs.

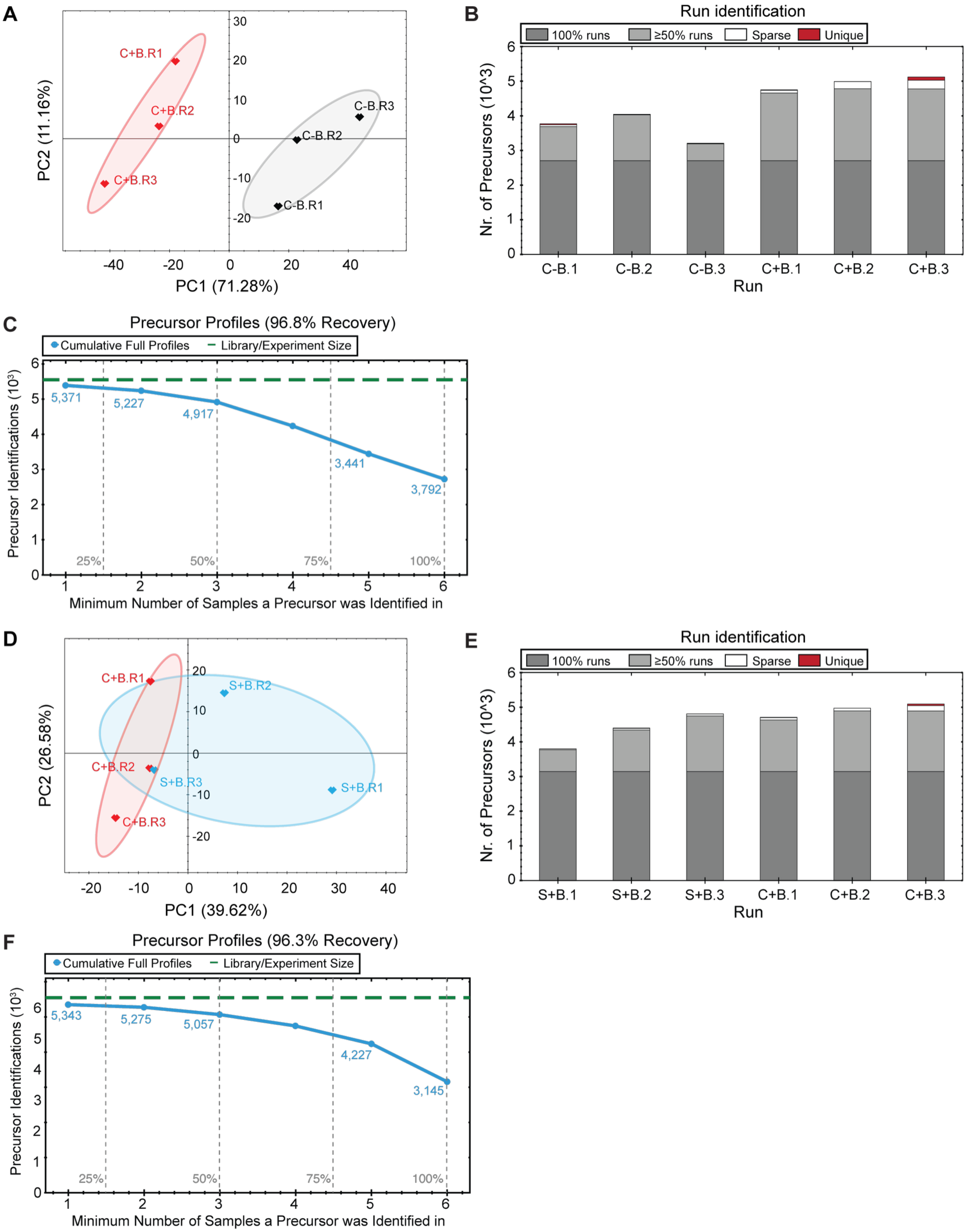


**Figure S14. Additional metrics for mPFC Nrxn-TurboID proteomics after acute cocaine exposure. (A)** Principal component analysis of mPFC Nrxn-TurboID proteomics data shows separated clustering between experimental samples (Cocaine+Biotin, red) and control samples

(Cocaine-Biotin, gray). **(B)** Number of precursors identified in 100% of runs (dark gray), in at least 50% of runs (light gray), in more than 1 but less than 50% of runs (white), or only in a single run (red). **(C)** Cumulative number of precursors identified per minimum number of runs a precursor was identified in. **(D-F)** Same as panels (A-C), except comparing Cocaine+Biotin vs Saline+Biotin.

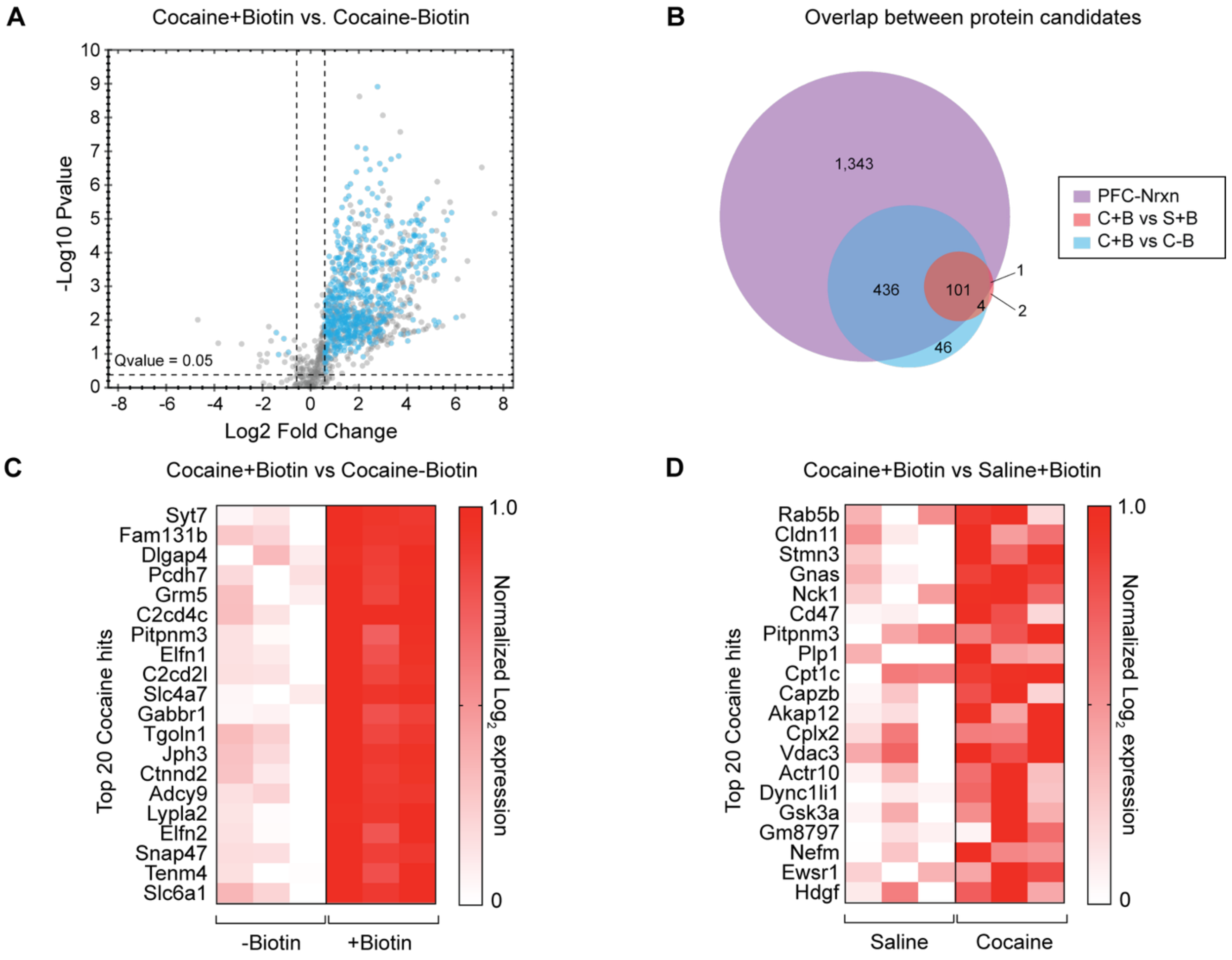


**Figure S15. Additional proteomics analysis after acute cocaine exposure. (A)** Volcano plot showing proteins enriched from mPFC cell bodies expressing Nrxn-TurboID in mice injected with Cocaine+Biotin vs Cocaine-Biotin. **(B)** Venn Diagram showing overlapping filtered Nrxn-TurboID candidates from the following datasets: mPFC-Nrxn (Figure 3), Cocaine+Biotin vs Saline+Biotin, Cocaine-Biotin vs Cocaine-Biotin. **(C)** Heatmap of min-max normalized $\log_2$ quantity of top 20 Nrxn-TurboID candidates comparing Cocaine+Biotin vs Cocaine-Biotin. **(D)** Heatmap of min-max normalized $\log_2$ quantity of top 20 Nrxn-TurboID candidates comparing Cocaine+Biotin vs Saline+Biotin.
